# Small-Molecule *In Vitro* Inhibitors of the Coronavirus Spike – ACE2 Protein-Protein Interaction as Blockers of Viral Attachment and Entry for SARS-CoV-2

**DOI:** 10.1101/2020.10.22.351056

**Authors:** Damir Bojadzic, Oscar Alcazar, Jinshui Chen, Peter Buchwald

**Affiliations:** Diabetes Research Institute, Miller School of Medicine, University of Miami, Miami, Florida, USA; Department of Molecular and Cellular Pharmacology, Miller School of Medicine, University of Miami, Miami, Florida, USA

**Keywords:** ACE2, antiviral, coronavirus, COVID-19, hydroxychloroquine, protein-protein interaction, SARS-CoV-2, small-molecule inhibitors, spike protein

## Abstract

Inhibitors of the protein-protein interaction (PPI) between the SARS-CoV-2 spike protein and ACE2, which acts as a ligand-receptor pair that initiates the viral attachment and cellular entry of this coronavirus causing the ongoing COVID-19 pandemic, are of considerable interest as potential antiviral agents. While blockade of such PPIs with small molecules is more challenging than with antibodies, small-molecule inhibitors (SMIs) might offer alternatives that are less strain- and mutation-sensitive, suitable for oral or inhaled administration, and more controllable / less immunogenic. Here, we report the identification of SMIs of this PPI by screening our compound-library that is focused on the chemical space of organic dyes. Among promising candidates identified, several dyes (Congo red, direct violet 1, Evans blue) and novel drug-like compounds (DRI-C23041, DRI-C91005) inhibited the interaction of hACE2 with the spike proteins of SARS-CoV-2 as well as SARS-CoV with low micromolar activity in our cell-free ELISA-type assays (IC_50_s of 0.2-3.0 μM); whereas, control compounds, such as sunset yellow FCF, chloroquine, and suramin, showed no activity. Protein thermal shift assays indicated that the SMIs identified here bind SARS-CoV-2-S and not ACE2. Selected promising compounds inhibited the entry of a SARS-CoV-2-S expressing pseudovirus into ACE2-expressing cells in concentration-dependent manner with low micromolar IC_50_s (6-30 μM). This provides proof-of-principle evidence for the feasibility of small-molecule inhibition of PPIs critical for coronavirus attachment/entry and serves as a first guide in the search for SMI-based alternative antiviral therapies for the prevention and treatment of diseases caused by coronaviruses in general and COVID-19 in particular.

## INTRODUCTION

COVID-19, which reached pandemic levels in early 2020 (WHO; March 11, 2020), is caused by the severe acute respiratory syndrome-coronavirus 2 (SARS-CoV-2) (*1–3*). SARS-CoV-2 is the most infectious agent in a century (*4*), having already caused more than forty million infections and a million deaths worldwide. This coronavirus (CoV) is an enveloped, positive-sense RNA virus with a large RNA genome of roughly 29.9 kilobases and a diameter of up to about 120 nm, characterized by club-like spikes emerging from its surface (*5, 6*). It is the most recently emerged among the seven CoVs known to infect humans. They include four CoVs that are responsible for about a third of the common cold cases (HCoV 229E, OC43, NL63, and HKU1) and three that caused epidemics in the last two decades associated with considerable mortality: SARS-CoV-1 (2002–2003, ~10% mortality), MERS-CoV (Middle East respiratory syndrome coronavirus; 2012, ~35% mortality), and now SARS-CoV-2 (2019–2020), which seems to be less lethal but more transmissible (*7, 8*). While the SARS-CoV-2 situation is still evolving, current estimates indicate that about 3% of infected individuals need hospitalization and the average infection fatality ratio (IFR, percentage of those infected that do not survive) is around 0.5% but in a strongly age-dependent manner, i.e., increasing from 0.001% in <20 years old to 8.3% in those >80 years old (*9*) (to be compared with an IFR of <0.1% for influenza). This created unprecedented health and economic damage and a correspondingly significant therapeutic need for possible preventive and/or curative treatments. As future CoVs that are highly contagious and/or lethal are also likely to emerge, novel therapies that could neutralize multiple strains are of particular interest especially as the large WHO Solidarity trial suggests that repurposed antiviral drugs including hydroxychloroquine, remdesivir, lopinavir, and interferon-β1, appear to have little or no effect on hospitalized COVID-19 patients, as indicated by overall mortality, initiation of ventilation, and duration of hospital stay (*10*).

Viral attachment and entry are of particular interest among possible therapeutic targets in the life cycle of viruses (*7*) because they represent the first steps in the replication cycle and take place at a relatively accessible extracellular site; they have indeed been explored for different viruses (*11*). CoVs use the receptor-binding domain (RBD) of their glycosylated S protein to bind to cell specific surface receptors and initiate membrane fusion and virus entry. For both SARS-CoV and SARS-CoV-2, this involves binding to angiotensin converting enzyme 2 (ACE2) followed by proteolytic activation by human proteases (*3, 5, 12, 13*). Hence, blockade of the RBD– hACE2 protein-protein interaction (PPI) can disrupt infection efficiency, and most vaccines and neutralizing antibodies (nAbs) aim to abrogate this interaction (*14, 15*). CoV nAbs, including those identified so far for SARS-CoV-2, primarily target the trimeric S glycoproteins, and their majority recognizes epitopes within the RBD that binds the ACE2 receptor (*15–19*). It would be important to have broadly cross-reactive nAbs that can neutralize a wide range of viruses that share similar pathogenic outcomes (*17*). The S proteins of SARS‐CoV, MERS‐CoV, and SARS-CoV‐2 have similar structures with 1100–1300 amino acids and RBDs spanning about 200 residues and consisting of core and external subdomains, with the RBD cores being responsible for the formation of S trimers – similarities that allow the possibility of broad neutralization (*20, 21*). SARS-CoV and SARS-CoV-2 share ~80% amino acid identity in their S proteins (*15, 20*); nevertheless, most current evidence indicates that SARS-CoV antibodies are not cross-reactive for SARS-CoV-2 (*22*). For example, one study found that none of the 206 RBD-specific monoclonal antibodies derived from single B cells of eight SARS-CoV-2 infected individuals cross-reacted with SARS-CoV or MERS-CoV RBDs (*23*). Antibody-like monobodies designed to bound to the SARS-CoV-2 S protein also did not bind that of SARS-CoV (*24*). As a further complicating factor, RNA viruses accumulate mutations over time, which yields antibody resistance and requires the use of antibody cocktails to avoid mutational escape (*25*).

In addition to being too highly target-specific, antibodies, as all protein therapies, are hindered by problems related to their solubility, unsuitability for oral or inhaled administration, and immunogenicity. By being foreign proteins, they themselves can act as antigens and elicit strong immune response in certain patients (*26–28*), and this is only further exacerbated by their long elimination half-lives (*29*). Even among FDA approved therapeutics, there were more post-market safety issues with biologics than with small-molecule drugs (*30*). Hence, peptides or small molecules can offer alternative approaches. Some peptide disruptors of this PPI have also been reported, but so far none have been very effective (*22, 31–33*). More importantly, because of bioavailability, metabolic instability (short half-life), lack of membrane permeability, and other issues, developing peptides into clinically approved drugs is difficult and rarely pursued (*34, 35*).

Small molecules traditionally were not considered for PPI modulation because they were deemed unlikely to be successful due to the lack of well-defined binding pockets on the protein surface that would allow their adequate binding. During the last decade, however, it has become increasingly clear that SMIs can be effective against certain PPIs. There are now >40 PPIs targeted by SMIs that are in preclinical development (*36–42*), and two of them (venetoclax (*43*), lifitegrast (*44*)) were recently approved by the FDA for clinical use (*45, 46*). Notably, the success of two small-molecule drugs that target HIV-1 entry and are now approved for clinical use, enfuvirtide and maraviroc, validates this strategy of antiviral drug discovery. Maraviroc targets the C-C motif chemokine receptor 5 (CCR5), a host protein used as a co-receptor during HIV-1 entry, and it is a noncompetitive allosteric inhibitor that stabilizes a conformation no longer recognized by the viral envelope (*11, 47*). Hence, it is an allosteric SMI of a PPI, highlighting the feasibility of such an approach to prevent viral entry. Interestingly, maraviroc has been claimed recently to inhibit the SARS-CoV-2 S-protein mediated cell fusion in cell culture (*6*). Therefore, SMIs could yield antiviral therapies that are more broadly active (i.e., less strain- and mutation-sensitive), more patient friendly (i.e., suitable for oral or inhaled administration), less immunogenic, and more controllable (shorter half-life / better biodistribution) than antibodies (*48*). Oral bioavailability offers a major advantage for access, wide-spread usage, and compliance (*49*) a requisite for long-term and broadly acceptable preventive use (*50–52*). For COVID-19, the possibility of direct delivery into the respiratory system via inhaled or intranasal administration is also important and unlikely to be achievable for antibodies. Broadly specific activity could make possible multi-strain or even pan-CoV inhibition, and while it is unlikely with antibodies (*22, 23*), it is possible for SMIs. For example, we have shown that while the corresponding antibodies did not cross-react for the human vs mouse CD40–CD40L PPI, our SMIs did so and had about similar potencies (*53, 54*).

Since previously we found that starting from organic dyes one can identify SMIs for co-signaling PPIs as potential immunomodulatory agents (*48, 53–59*), we initiated a screen of such compounds for their ability to inhibit the SARS-CoV-2-S–ACE2 PPI. This led to the identification of several organic dyes (**1**–**5**, Figure 1) that show promising inhibitory activity of this PPI *in vitro*, including methylene blue (**6**), a phenothiazine dye approved by the FDA for the treatment of methemoglobinemia, which we have described separately (*60*). More importantly, it also led to the identification of new and more potent SMIs (**7**–**13**) that are more drug-like and no longer contain color-causing chromophores as summarized below.

**Figure 1.**
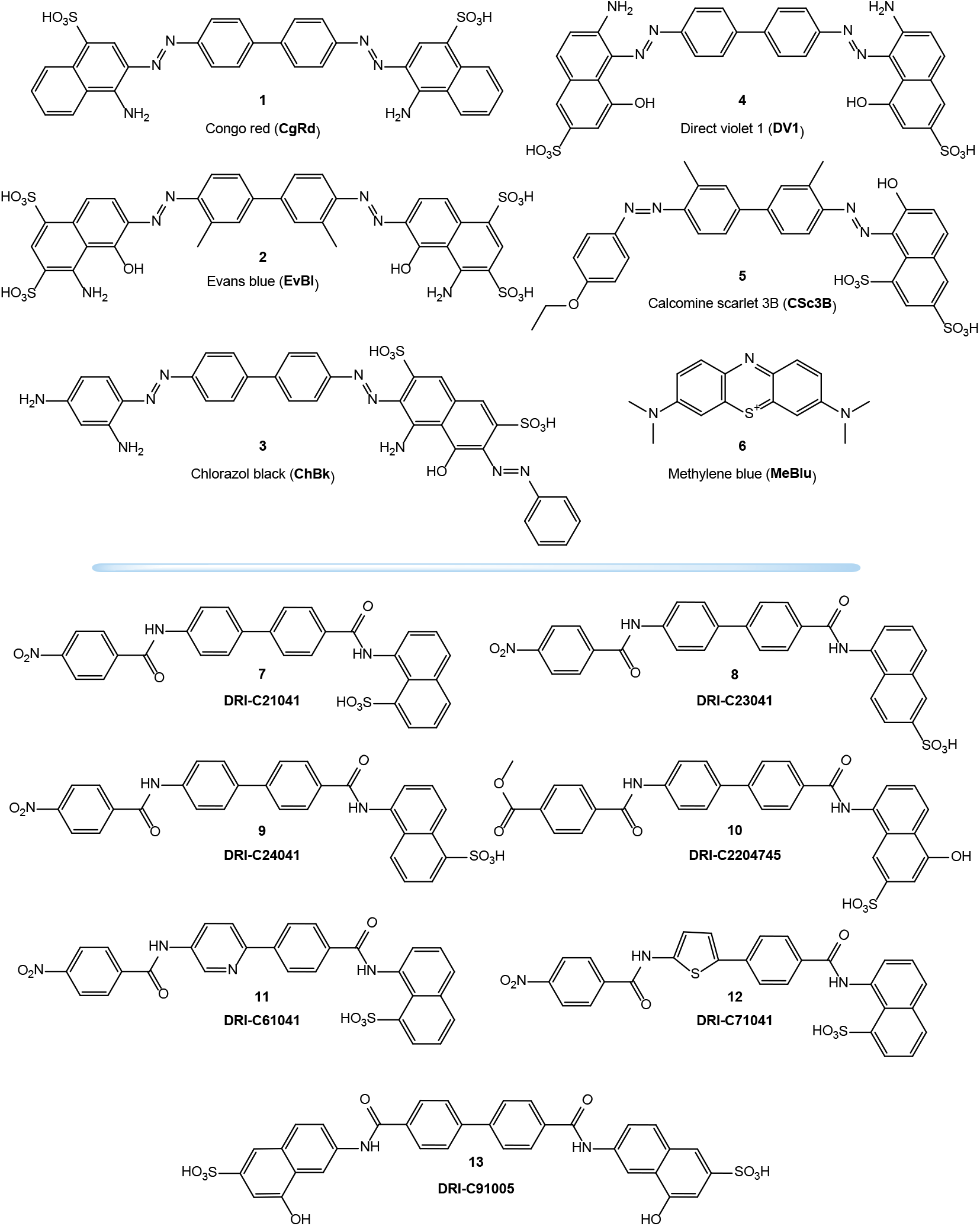
Compounds of the present study. Chemical structures of the organic dye (**1**–**6**) and non-dye DRI-C compounds (**7**–**13**) used in the present study.

## RESULTS

As part of our work to identify SMIs for co-signaling PPIs that are essential for the activation and control of immune cells, we discovered that the chemical space of organic dyes, which is particularly rich in strong protein binders, offers a useful starting point. Accordingly, it seemed logical to explore it for possible inhibitors of the SARS-CoV-2 S protein – ACE2 PPI that is an essential first step for the viral entry of this novel, highly infectious coronavirus. We were able to set up a cell-free ELISA-type assay to quantify the binding of SARS-CoV-2 S protein (as well as its SARS-CoV analog) to their cognate receptors (human ACE2) and used this to screen our existing in-house compound library containing a large variety of organic dyes and a set of colorless analogs prepared as potential SMIs for costimulatory PPIs. These maintain the main molecular framework of dyes but lack the aromatic azo chromophores responsible for the color as they are replaced with amide linkers (*58, 59*).

### Chemistry and Synthesis

All new compounds used here were synthesized as described before as part of our effort to identify novel SMIs for the CD40–CD40L costimulatory PPI (*58, 59*). Synthesis involved one or two amide couplings (using a modified version of the procedure from (*61*)) and a hydrogenation (using a modified version of the procedure from (*62*)). These steps were used with different linkers and naphthyl moieties as needed for each structure; all corresponding details are summarized in the Supplementary Material (Supplementary Methods and Supplementary Schemes S1-S6). All structures tested here (**1**–**13**) are shown in Figure 1.

### Screening Assays

As a first step, we explored the feasibility of setting up screening assays using a cell-free ELISA-type format similar to those used in our previous works with Fc-conjugated receptors coated on the plate and FLAG- or His-tagged ligands in the solution (*53, 56–58*). Concentration-response assessments of binding to ACE2 indicated that both the S1 and RBD portions of SARS-CoV-2-S bind strongly and follow classic sigmoid patterns corresponding to the law of mass action (*63*) with a slightly stronger binding for RBD than S1 (Figure 2). Fitting of data gave median effective concentrations (EC_50_s) and hence binding affinity constant (*K*_d_) estimates of 3.7 and 14.7 nM, respectively (98 and 1125 ng/mL) – in good agreement with the specifications of the manufacturer (SinoBiological; Wayne, PA, USA) and published values indicating a low nanomolar range (4–90 nM) typically based on surface plasmon resonance (SPR) studies (*5*). Because we are interested in possible broad-spectrum inhibitors, we also performed concentration-response assessments of the binding of SARS-CoV and HCoV-NL63 S proteins (using their S1&S2 and S1 domains, respectively) as they also use ACE2 as their cognate receptor. SARS-CoV bound with about similar potency as SARS-CoV-2 (13.9 nM; 1843 ng/mL), whereas HCoV-NL63 had significantly lower affinity (45.8 nM, 3610 ng/mL) (Figure 2).

**Figure 2.**
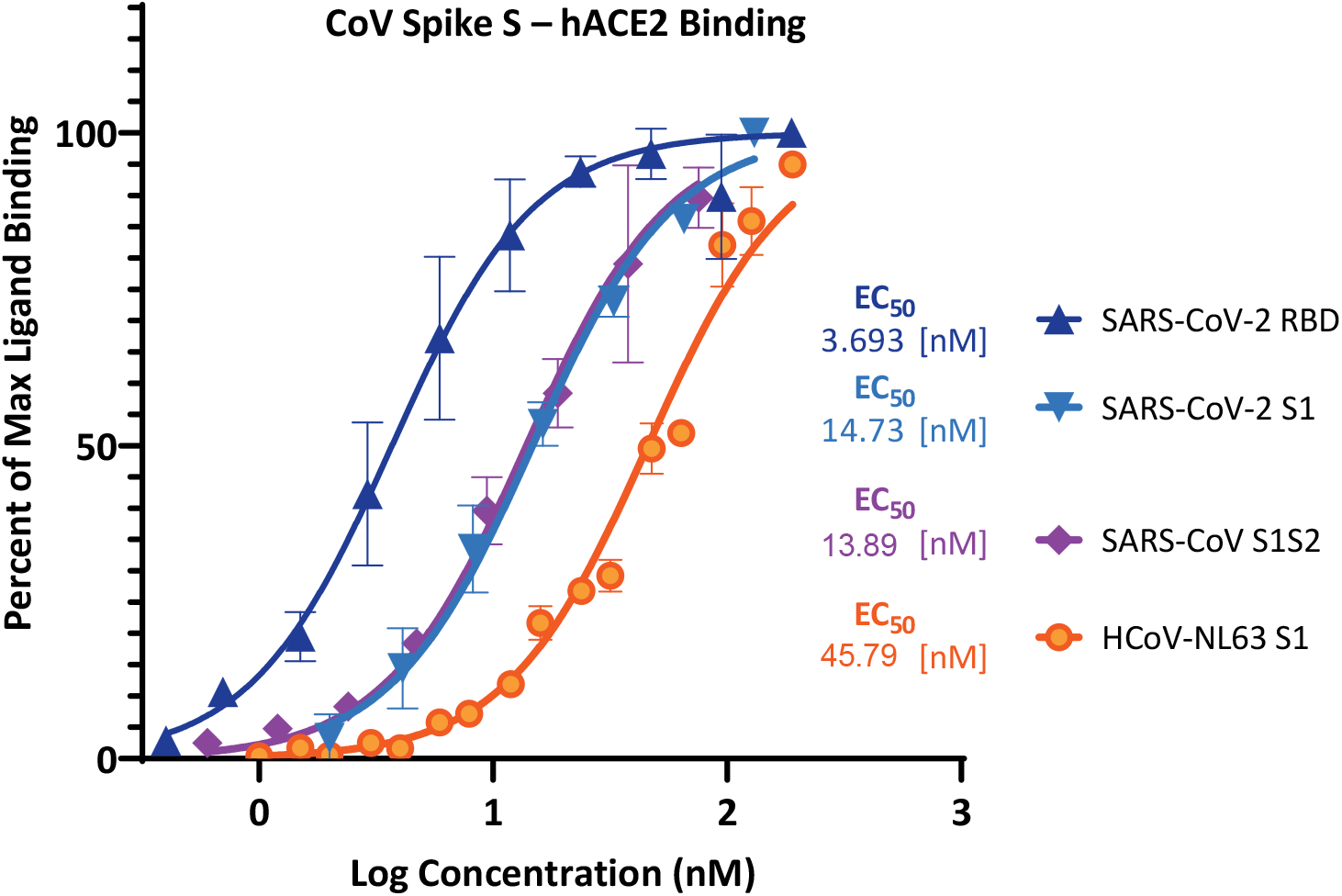
Concentration-response curves for binding of CoV spike protein domains to human ACE2 in cell-free ELISA-type assays. Binding curves and corresponding EC_50_s are shown for SARS-CoV-2 (RBD and S1), SARS-CoV (S1&S2), and HCoV-NL63 (S1). They were obtained using Fc-conjugated hACE2 coated on the plate and His-tagged S1, S1S2, or RBD added in increasing amounts as shown with the amount bound detected using an anti-His–HRP conjugate (mean ± SD for two experiments in duplicates).

Based on this, we first used this assay to screen for inhibitors of SARS-CoV-2 RBD binding, which showed the strongest affinity to hACE2. In fact, this assay setup is very similar to one recently shown to work as a specific and sensitive SARS-CoV-2 surrogate virus neutralization test based on antibody-mediated blockage of this same PPI (CoV-S–ACE2) (*64*). We screened our in-house library of organic dyes plus existing analogs together with a few additional compounds that are or have been considered of possible interest in inhibiting SAR-CoV-2 by different mechanisms of action, e.g., chloroquine, clemastine, and suramin (*22, 65–68*). Screening at 5 μM indicated that most have no activity and, hence, are unlikely to interfere with the S-protein – ACE2 binding needed for viral attachment. Nevertheless, some showed activity (Supplementary Material, Figure S1). Compounds showing the strongest activity, i.e., rose Bengal, erythrosine B (ErB), and phloxine B, are known promiscuous SMIs of PPIs (*56*). As such, they are of no value here being nonspecific; they were included as positive controls. This screening also identified methylene blue (MeBlu, **6**), a phenothiazine dye approved by the FDA for the treatment of methemoglobinemia and also used for several other therapeutic applications in the developed world (*69–71*) and with additional potential for certain developing world applications such as malaria (*72*), as showing promising inhibitory activity for the SARS-CoV-2-S–hACE2 PPI, likely contributing to its anti-CoV activity (*73*); this has been discussed separately (*60*).

### Binding Inhibition (Concentration-Response)

Next, detailed concentration-response assessments were performed to establish inhibitory activity (IC_50_) per standard experimental guidelines in pharmacology and experimental biology (*74, 75*). These confirmed that indeed several organic dyes as well as non-dye DRI compounds inhibited this PPI in a concentration-dependent manner with low micromolar IC_50_s (Figure 3). For example, among tested dyes, Congo red (CgRd, **1**), direct violet 1 (DV1, **4**), Evans blue (EvBl, **2**), chlorazol black (ChBk, **3**), and calcomine scarlet 3B (CSc3B, **5**) had IC_50_s of 0.99, 1.44, 2.25, 2.57, and 4.25 μM, respectively. Further, we also found several DRI compounds of low micromolar activity including some, such as DRI-C91005 (**13**) and DRI-C23041 (**8**), with even better submicromolar IC_50_s (160 and 520 nM, respectively). For the compounds tested here, concentration dependencies were adequately described by a standard log inhibitor vs response model (i.e., a classical sigmoid binding function with a Hill slope of 1 (*63*)). Sunset yellow FCF (FD&C yellow #6), an azo dye and an FDA approved food colorant included as a possible negative control, showed no inhibitory activity. Notably, neither chloroquine nor suramin showed inhibitory activity in this assay. We tested chloroquine, an anti-parasitic and immunosuppressive drug primarily used to prevent and treat malaria, because it was the subject of considerable controversy regarding its potential antiviral activity against SARS-CoV-2 (*65*). We also tested suramin, an antiparasitic drug approved for the prophylactic treatment of African sleeping sickness (trypanosomiasis) and river blindness (onchocerciasis), because it was claimed to inhibit SARS-CoV-2 infection in cell culture by preventing binding or entry of the virus (*68*) and because it was one of the first compounds we found to inhibit the CD40–CD40L PPI (*55*). On the other hand, erythrosine B (ErB, FD&C red #3), an FDA approved food colorant that we found earlier to be a promiscuous PPI inhibitor and have been using as positive control in such assays, inhibited with an IC_50_ of 0.4 μM, similar to its activity found for other PPIs tested before (1–20 μM) (*56*).

**Figure 3.**
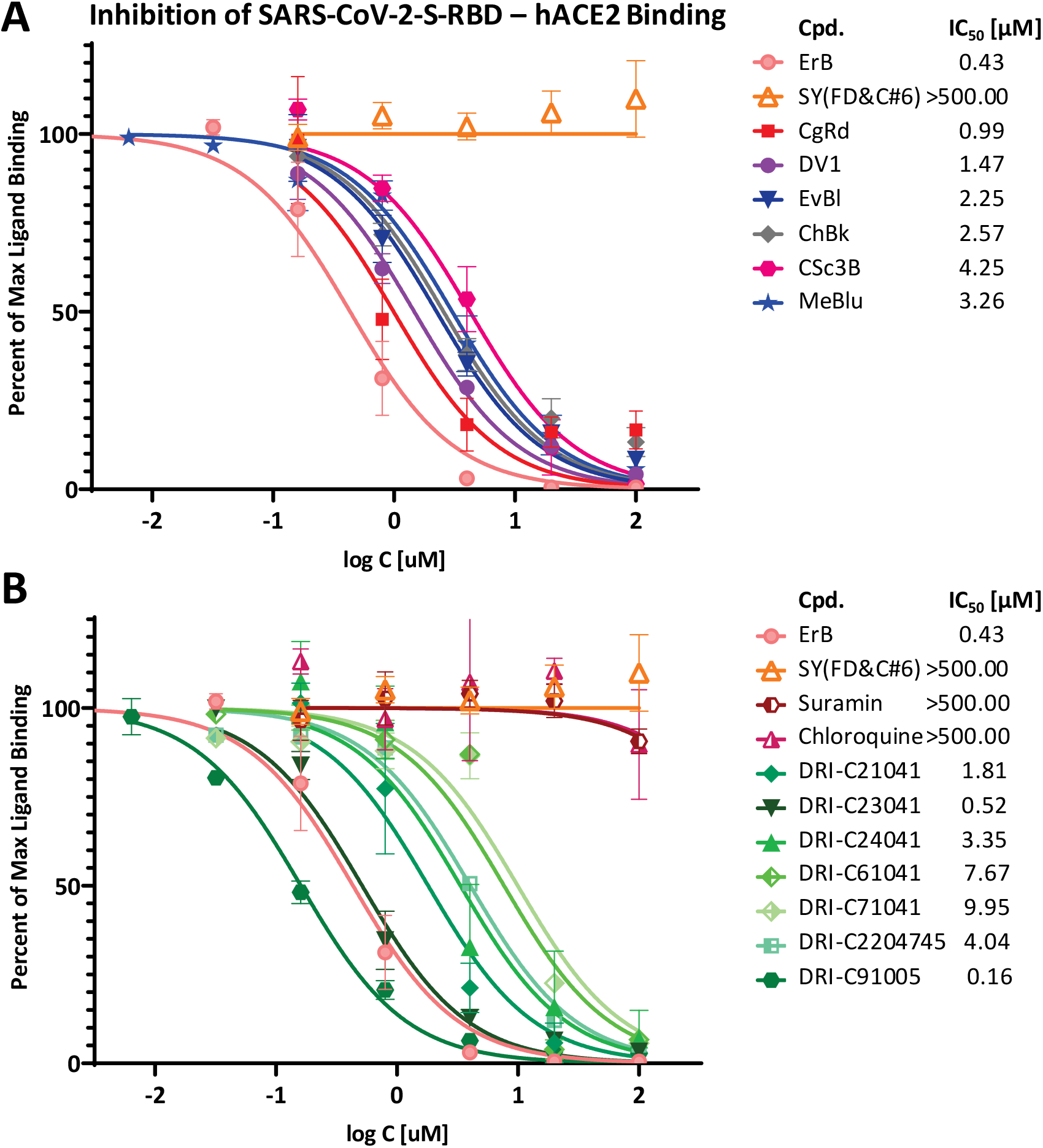
Concentration-dependent inhibition of SARS-CoV-2-S-RBD binding to ACE2 by compounds of the present study. Concentration-response curves obtained for the inhibition of the PPI between SARS-CoV-2-RBD (His-tagged, 0.5 μg/mL) and hACE2 (Fc-conjugated, 1 μg/mL) in cell-free ELISA-type assay with dye (**A**) and non-dye (**B)** compounds tested. The promiscuous PPI inhibitor erythrosine B (ErB) and the food colorant FD&C yellow no. 6 (sunset yellow, SY) were included as a positive and negative controls, respectively. Data are mean ± SD from two experiments in duplicates and were fitted with standard sigmoid curves for IC_50_ determination. Estimated IC_50_s are shown in the legend indicating that while suramin and chloroquine were completely inactive (IC_50_ > 500 μM), several of our in-house compounds including organic dyes (CgRd, DV1, and others) as well as proprietary DRI-C compounds (e.g., DRI-C23041, DRI-CC91005) showed promising activity, some even at sub-micromolar levels (IC_50_ < 1 μM).

For a few representative compounds, we also tested their ability to inhibit not just the binding of SARS-CoV-2-RBD but also that of SARS-CoV-2-S1 to hACE2. We obtained similar potencies; e.g., DRI-C23041 had an IC_50_ of 1.88 μM (95% CI of 1.32–2.68 μM) for S1 (Supplementary Material, Figure S2) vs 0.52 μM (95% CI of 0.42–0.63 μM) for RBD (Figure 3). This confirms that these are indeed real inhibitory activities relevant for the S protein – hACE2 PPI of interest. More importantly, we also assessed the ability of selected promising compounds to inhibit the binding of SARS-CoV-S to ACE2 using a similar setup. As shown in Figure 4, several of the same compounds including organic dyes (CgRd, DV1, and others) as well as DRI compounds showed similar activity against SARS-CoV as against SARS-CoV-2. For compounds tested in this assay such as CgRd, CVN, EvBl, CSc3B, DRI-C23041, and DRI-C91005 the IC_50_s were 3.9, 2.6, 1.3, 9.9, 3.4, and 0.24 μM (Figure 4), respectively – values that are similar to those obtained for SARS-CoV-2 inhibition (Figure 3), raising the possibility of broad-spectrum anti-CoV activity.

**Figure 4.**
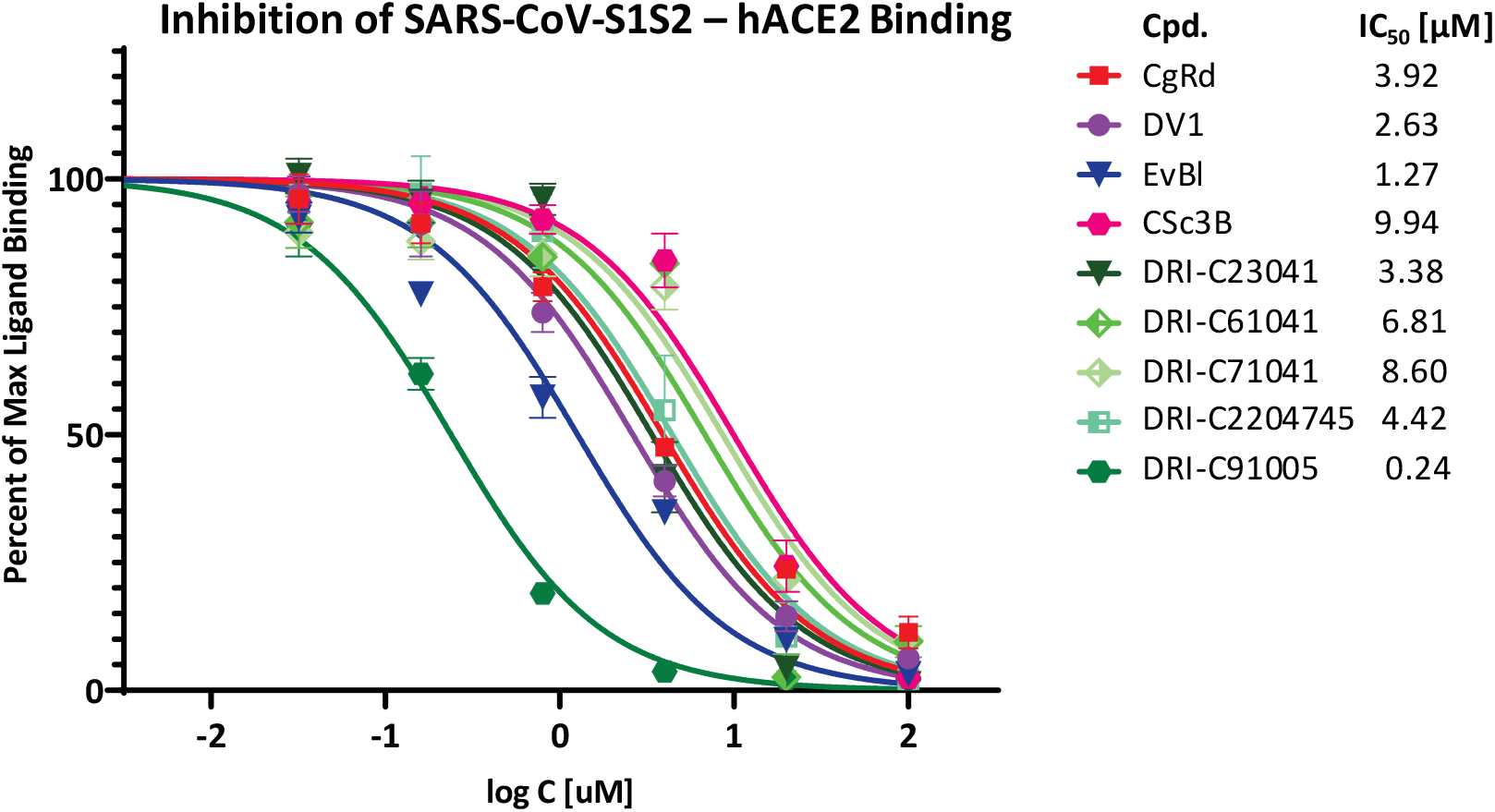
Concentration-dependent inhibition of SARS-CoV-S1S2 binding to ACE2 by representative compounds of the present study. Concentration-response curves obtained for the inhibition of the PPI between SARS-CoV-S1S2 (His-tagged, 1 μg/mL) and hACE2 (Fc-conjugated, 1 μg/mL) in cell-free ELISA-type assay by selected representative dye and non-dye compounds. Data and fit as before (Figure 3). Most compounds including several DRI-C compounds show similar activity against SARS-CoV (i.e., SARS-CoV-1) as against SARS-CoV-2 raising the possibility of broad-spectrum activity.

Besides activity, it is also important to achieve adequate selectivity, specificity, and safety. To become promising lead candidates, small-molecule compounds are usually expected to show >30-fold selectivity over other possible pharmacological targets of interest (*76, 77*). As a counter-assay, here we assessed inhibitory activity against the TNF-R1–TNF-α interaction, as we have done before (*58, 59*). Most of the dyes found here to inhibit the SARS-CoV-2–ACE2 PPI (Figure 3) seem to be relatively promiscuous as they also inhibited the TNF-R1–TNF-α PPI (Figure 5) showing only some limited selectivity, e.g., 6-fold for CgRd (0.99 vs 6.0 μM) as one of the best and only 1.4-fold for DV1 (1.5 vs. 2.1 μM). On the other hand, several DRI-C compounds showed good, more than 100-fold selectivity, e.g., >400-fold for DRI-C23041 (0.52 vs 233 μM). The symmetric DRI-C91005 seems an exception that was the most potent in all assays but showed no selectivity (0.16 vs 0.16 μM). As these DRI-C compounds were designed to target CD40–CD40L, they all inhibit that PPI with high nanomolar – low micromolar potency (see Discussion).

**Figure 5.**
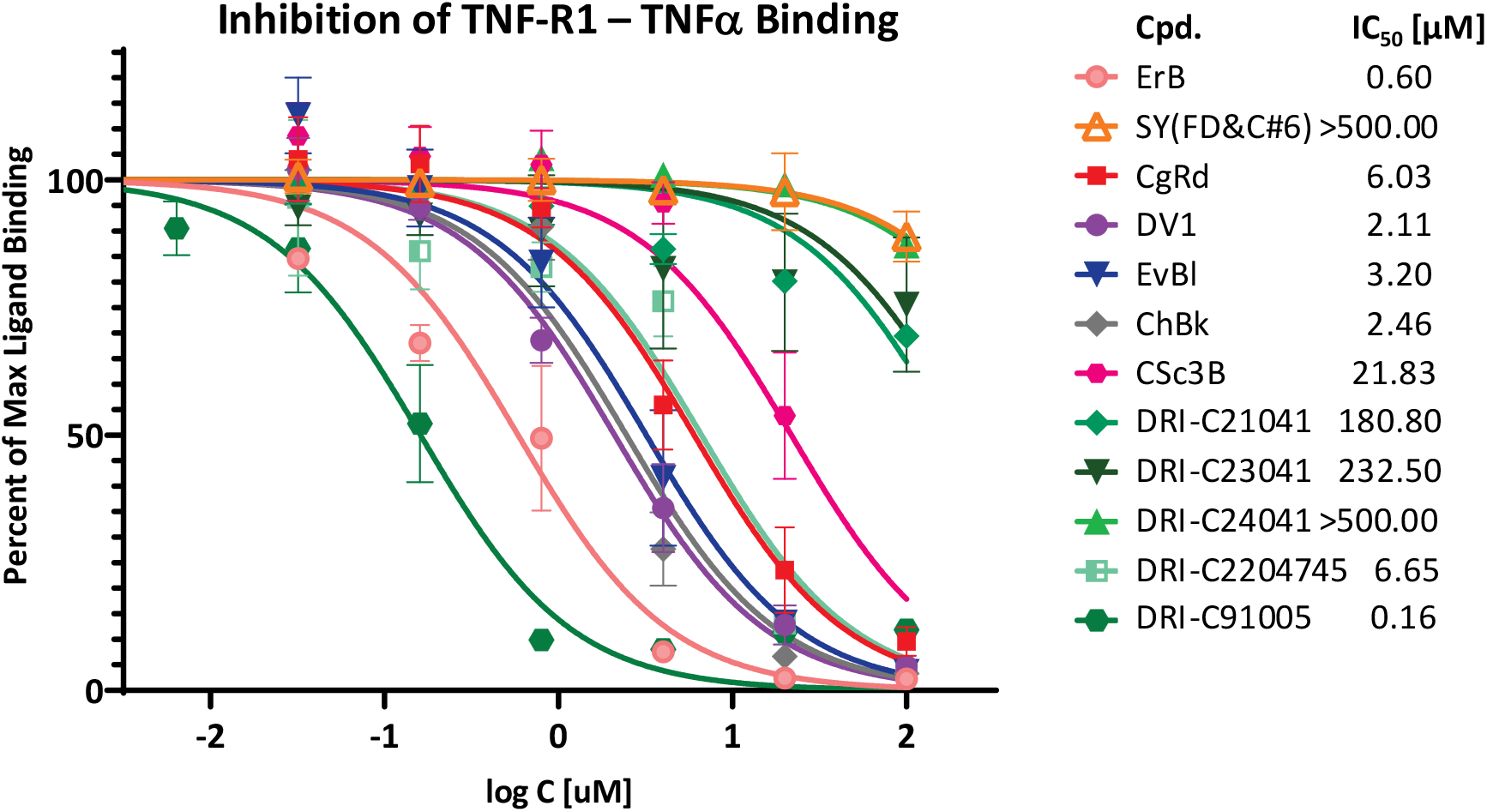
Concentration-dependent inhibition of TNF-R1–TNFα by compounds of the present study. Concentration-response curves obtained for the inhibition of this important TNF superfamily PPI in similar cell-free ELISA-type assay as used for the CoV-S–ACE2 PPIs to assess selectivity. Data and fit as before (Figure 3). As the IC_50_ values indicate, DRI-C compounds showed more than 100-fold selectivity in inhibiting the CoV-S PPI vs the TNF PPI.

### Binding Partner (Protein Thermal Shift)

To establish whether these SMIs bind to CoV-S or ACE2, we used a protein thermal shift assay (differential scanning fluorimetry or ThermoFluor assay) (*78, 79*) as we did before for CD40L (*59*). This assay quantifies the shift in protein stability caused by binding of a ligand via use of a dye whose fluorescence increases when exposed to hydrophobic surfaces, which happens as the protein starts to unfold as it is heated and exposes its normally buried hydrophobic core residues. It allows rapid and inexpensive evaluations of the temperature-dependence of protein stability using real-time PCR instruments and only small amounts of protein. It is sensitive enough to assess small-molecule PPI interference and can be used even as a screening assay (*80*). As shown in Figure 6, the presence of CgRd or DRI-C23041 caused clear left-shifts in the melting temperature (*T*_m_) of the protein for SARS-CoV-2-RBD, but not ACE2 (purple vs. blue lines) indicating the former as the binding partner. This is encouraging, as SMIs targeting the S-protein are much more likely to (1) not cause undesirable side effects than ACE2-targeting ones, which could interfere with ACE2 signaling, and (2) be more broadly specific due to the structural similarity of the different CoV S glycoproteins. Binding of a ligand usually results in an increase (right-shift) of the melting temperature due to stabilization of the protein; however, cases with a decrease (hence, destabilization) have also been reported (*81*) including for the Ebola virus glycoprotein (*82*).

**Figure 6.**
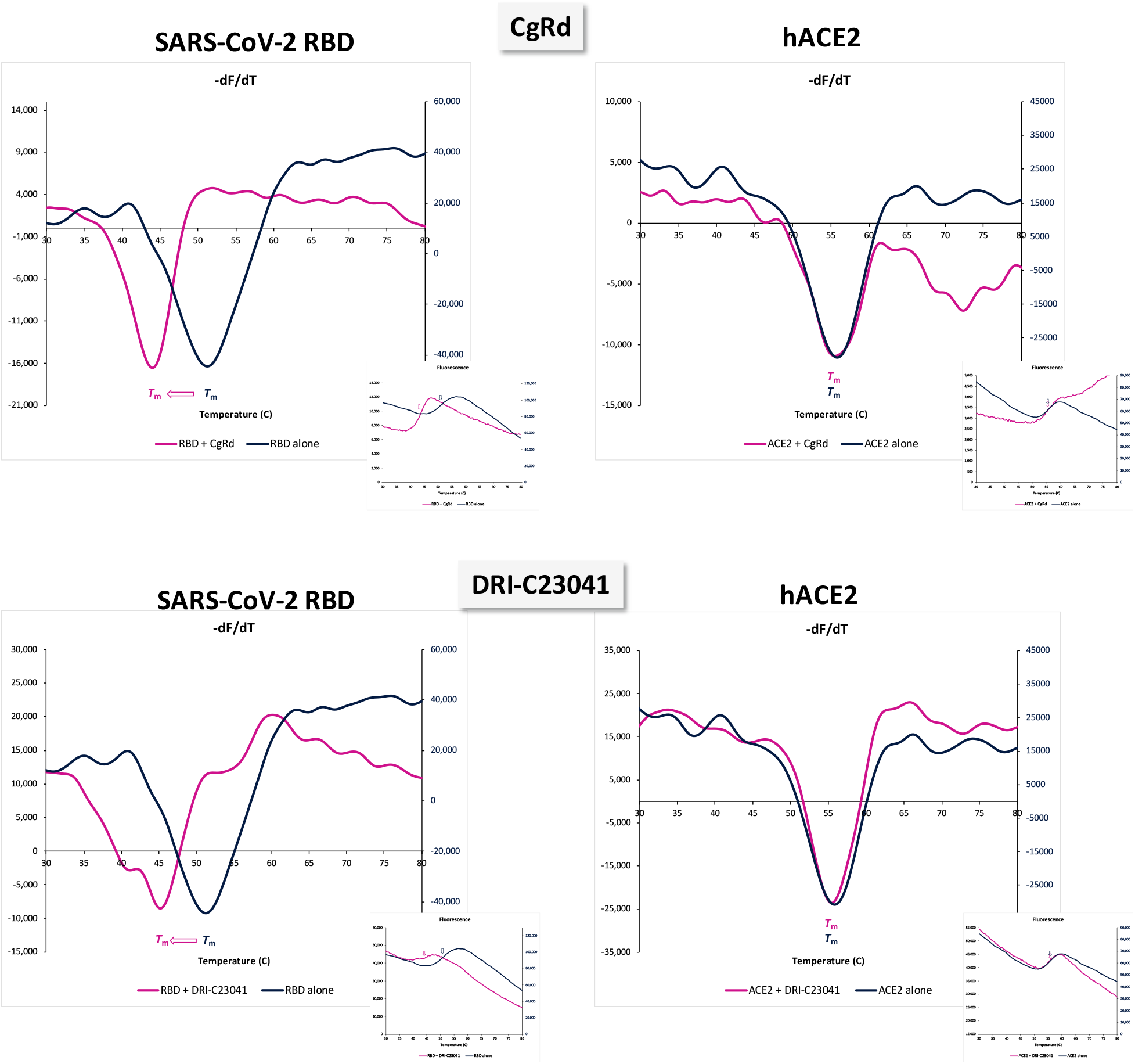
Identification of the binding partner by protein thermal shift. Differential scanning fluorimetry assay indicating SARS-CoV-2 RBD and not ACE2 as the binding partner of the present SMI compounds. The presence of Congo red (top) or DRI-C23041 (bottom) at 10 μM caused clear shifts in the melting temperature of the protein for RBD as indicated by the derivatives d*F*/d*T* (left; purple vs. blue line), but not for hACE2 (right) (smaller insets are normalized fluorescence *F* data).

### Inhibition of SARS-CoV-2 Pseudo-Virus Entry

For a set of selected active compounds, we were able to confirm that they also inhibit viral entry. This has been done with pseudoviruses bearing the SARS-CoV-2 S spike protein (plus fluorescent reporters) and generated using BacMam-based tools. These allow quantification of viral entry, as they express bright green fluorescent protein that is targeted to the nucleus of ACE2- (and red fluorescence reporter) expressing host cells (here, HEK293T), but can be handled using biosafety level 1 containment, as they do not replicate in human cells. A day after entry, host cells express green fluorescence in the nucleus, indicating pseudovirus entry. If entry is blocked, the cell nucleus remains dark. In this assay, several of our SMIs tested, for example, CgRd, DV1, and DRI-C23041 showed good concentration-dependent inhibition as illustrated by the corresponding images and bar graphs in Figure 7. Fitting with regular concentration response curves indicated a very encouraging IC_50_ of 5.8 μM for DRI-C23041. CgRd and DV1 also inhibited, but with higher IC_50_s (26 and 64 μM for, respectively), which is not unexpected for such azo dyes as they tend to lose activity in cell-based assay due to nonspecific binding (Figure 7C).

**Figure 7.**
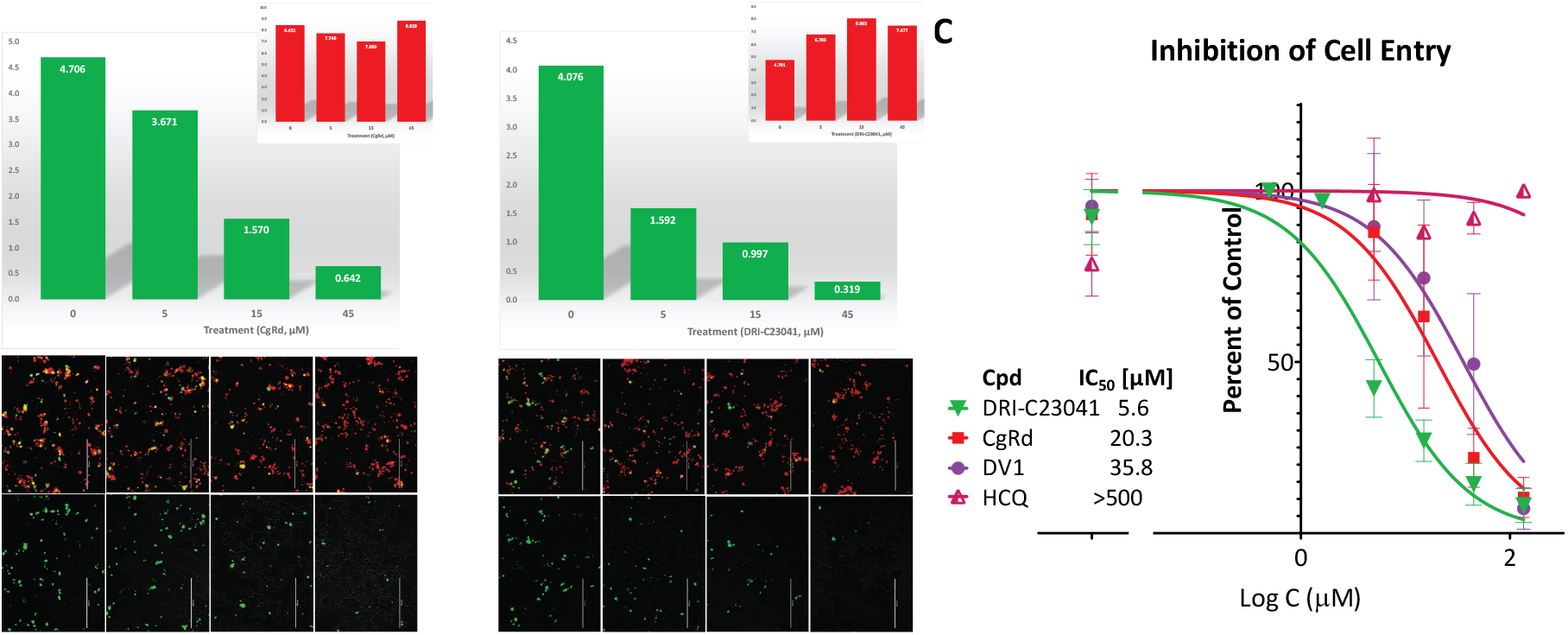
Concentration-dependent inhibition of SARS-CoV-2 pseudovirus entry into hACE2 expressing host cells by selected compounds. Quantification of entry of pseudoviruses bearing the SARS-CoV-2 S protein (plus green fluorescent protein reporters; BacMam-based) in ACE2 (plus red fluorescence) expressing host cells (HEK293T). Representative images (bottom row) and their quantification for pseudovirus (green) and ACE2 expression (red) using ImageJ (top row) are shown from one experiment for CgRd and DRI-C23041 in **A** and **B**, respectively; average data from three experiments fitted with typical concentration-response curves are shown in **C**. The amount of green present is proportional with the number of infected cells as green fluorescence is expressed only in pseudovirus infected cells, while amount of red is proportional with the number of ACE2-expressing cells. The organic dye CgRd (**A**), but especially DRI-C23041 (**B**) showed concentration-dependent inhibition with activities corresponding to low micromolar IC_50_ values, whereas hydroxychloroquine (HCQ) showed no effect (**C**).

As a first safety assessment, in parallel with the cell assays, we also evaluated cytotoxicity for several compounds in the same cells and at the same concentrations using a standard MTS assay to ensure that effects are present at non-toxic concentration levels. Notably, chloroquine already showed noticeable cytotoxicity at 45 μM concentrations in this assay with HEK293T cells, so its effect on pseudovirus entry could not be reliably evaluated there and hydroxychloroquine was used. We have shown before that compounds such as DRI-C21041 (**7**) or DRI-C24041 (**9**) did not have significant effects on the viability of THP-1 human cells for concentrations of up to 200 μM (*58, 59*). In line with that, DRI-C23041 (**8**) was the least cytotoxic among tested compounds here and showed no significant effects on HEK293T at 45 μM (Supplementary Material, Figure S3), whereas it had a strong effect on viral entry (Figure 7).

## DISCUSSION

Results obtained here confirm again that the chemical space of organic dyes can serve as a useful starting platform for the identification of SMI scaffolds for PPI inhibition. Organic dyes need to be good protein binders; hence, their contain privileged structures for protein binding (*83–85*) and can provide a better starting point toward the identification of SMIs of PPIs than most drug-like screening libraries, whose chemical space has been shown to not correspond well with that of promising PPI inhibitors (*86–88*). Using this strategy, we have identified promising SMIs for the CD40–CD40L costimulatory interaction (*53, 58, 59*) and even some promiscuous SMIs of PPIs (*56*). Of course, because most dyes are unsuitable for therapeutic applications due to their strong color and, in the case of azo dyes, their quick metabolic degradation (*89, 90*), structural modifications are needed to optimize their clinical potential (*58, 59*).

Here, we explored the potential of this approach to identify SMIs for the PPI between ACE2 and CoV spike proteins as potential antivirals inhibiting attachment. Since SARS-CoV-2 uses its S protein via its RBD to bind ACE2 as the first step of its entry (*3, 5, 12, 13*), targeting these proteins is a viable therapeutic strategy, and work with prior zoonotic CoV has demonstrated proof-of-concept validity for such approaches. By screening our compound library spanning the chemical space of organic dyes, we identified several promising SMIs including dyes, such as Congo red and direct violet 1, as well as novel drug-like compounds, such as DRI-C23041, that *(1)* inhibited the SARS-CoV-2-S–hACE2 PPI with low micromolar activity (Figure 3), *(2)* seem to bind to SARS-CoV-2-S and not ACE2 (Figure 6), and *(3)* inhibited entry of SARS-CoV-2-S displaying pseudoviruses into ACE2 expressing cells (Figure 7). Importantly, there is clear indication of a consensus structural motif present in the active compounds identified here: a biphenyl linker with a naphthyl at one end and another aromatic naphthyl or phenyl at the other end, both with at least one but preferably multiple polar substituents (Figure 1).

Following the emergence of SARS-CoV in the early 2000s, a limited number of groups performed high-throughput screening (HTS) assays to identify inhibitory drug candidates for targeting various early steps in its cell invasion. Identified candidates included some putative SMIs of viral entry, for example, SSAA09E2 (*91*) and VE607 (*92*). Inhibitory candidates acting by other mechanism identified included, for example, SSAA09E1, SSAA09E3 (*91*); MP576, HE602 (*92*); ARB 05-018137, ARB 05-090614 (*93*); KE22 (*94*); and others (reviewed in (*22, 31, 32*)). Most of these showed activity only in the low micromolar range, e.g., 3.1, 0.7, and 1.6 μM for SSAA09E2, K22, and VE607, respectively (*22*). Even if these compounds showed some evidence of inhibiting CoV infection, no approved preventive or curative therapy is currently available for human CoV diseases. In addition to the relatively low (micromolar) potency, a main reason for this is that these compounds were not suitable for clinical translatability. They could not pass the pre-clinical development stage and enter clinical trials due to their poor bioavailability, safety, and pharmacokinetics (*22*). Note that by starting from a different chemical space and not from that of drug-like molecules typically used for HTS, our best SMIs identified here are already well within this low micromolar range for SARS-CoV-2. There also was a recent attempt at identifying possible disruptors of the SARS-CoV-2-S-RBD–ACE2 binding using AlphaLISA assay based HTS of 3,384 small-molecule drugs and pre-clinical compounds suitable for repurposing that identified 25 possible hits (*95*). However, these were also of relatively low potency (micromolar IC_50_s). None of them shows resemblance with the scaffold(s) identified here – highlighting again the known lack of overlap between the chemical space of existing drugs / drug-like structures and that of PPI inhibitors.

The S protein is a homotrimer with each of its monomer units being about 180 kDa, and it contains two subunits, S1 and S2, mediating cell attachment and fusion of the viral and cellular membrane, respectively (*16, 96*). The RBD of the S protein is located within the S1 domain and is known to switch between a standing-up position for receptor binding and a lying-down position for immune evasion (*12, 31*). CoVs can utilize different receptors for binding, but several CoVs, even from different genera, can also utilize the same receptor. SARS-CoV-2 is actually the third human CoV utilizing ACE2 as its cell entry receptor, the other two being SARS-CoV and the α-coronavirus HCoV NL63 (*3*). MERS-CoV recognizes dipeptidyl peptidase 4 (DPP4) (*3–5*), while HCoV 229E recognizes CD13 (*97*). Some β-coronaviruses (e.g., HCoV OC43) bind to sialic acid receptors (*98*). Having access to broadly cross-reactive agents that can neutralize a wide range of antigenically disparate viruses that share similar pathogenic outcomes would be highly valuable from a therapeutic perspective (*17*), and SMIs are less specific and could yield therapies that are more broadly active (i.e., less strain- and mutation-sensitive) than antibodies, which tend to be highly specific. We have shown before that while the corresponding antibodies are species specific for the CD40–CD40L PPI, our SMIs could inhibit both the human and mouse system with similar potencies (*53, 54*). Hence, it is feasible that SMI structures can be identified that in addition to inhibiting SARS-CoV-2, also inhibit other CoVs, including the high lethality SARS-CoV and MERS-CoV as well as the common cold causing HCoVs. Along these lines, it is very encouraging that SMIs identified here target the CoV-S protein and not ACE2 (Figure 6) and they show similar potency in inhibiting SARS-CoV (Figure 4) and SARS-CoV-2 (Figure 3). Such inhibitory effects on viral attachment can translate into antiviral activity against SARS-CoV-2 and possibly other ACE2-binding CoVs such as SARS-CoV and the α-coronavirus HCoV NL63.

While the SMIs identified here are not very small structures (MW in the 550 to 700 Da range), they are still relatively small compared to typical SMIs of PPIs. These tend to have larger structures to achieve sufficient activity, and they often severely violate the widely used “rule-of-five” criteria, which, among others, requires MW < 500 (*99*). In the last two decades, this “rule” has been used as a guide to ensure oral bioavailability and an adequate pharmacokinetic profile. Nevertheless, an increasing number of new drugs have been launched recently (including the two small-molecule PPI inhibitors discussed earlier) that significantly violate these empirical rules proving that oral bioavailability can be achieved even in the “beyond rule-of-five” chemical space (*100*). Hence, our results provide further proof for the feasibility of SMI for CoV attachment and provide a first map of the chemical space needed to achieve this.

Finally, these DRI-C structures (**8**–**13**) were originally intended to modulate co-signaling interactions, specifically to inhibit the CD40–CD40L costimulatory interaction, and they do so with low micromolar potency in cell assays (≈10 μM) (*58, 59*). While some show good selectivity vs TNF (e.g., DRI-C23041, DRI-C24041), others seem more promiscuous (e.g., DRI-C91005). TNF-inhibitory activities here were somewhat stronger than those we obtained before, e.g., IC_50_s of 0.6 vs 5 (*56*) for ErB or 181 vs >1000 (*59*) for DRI-C21041, possibly due to the use of a different blocking buffer. We hope that these PPI inhibitory activities can be ultimately separated, but even if not and they still retain some activity in modulating co-signaling interactions, this might not necessarily be counterproductive. It could provide a unique opportunity to pursue dual-function molecules that on one hand, have antiviral activity by inhibiting the interaction needed for CoV attachment (e.g., SARS-CoV-2-S–ACE2) and, on the other, possess immunomodulatory activity to rein-in overt inflammation (inhibiting CD40–CD40L) or to unleash T cell cytotoxicity against virus-infected cells (inhibiting PD-1–PD-L1). Targeting of the PD-1 co-signaling pathway could be particularly valuable for its potential in restoring T cell homeostasis and function from an exhausted state (*101, 102*), which is of interest to improve viral clearance and rein-in the inflammatory immune response and the associated cytokine storm during anti-viral responses such as those likely implicated in the serious side effects seen in many COVID-19 patients (*1, 103–105*). Notably, the overexuberant immune response seen in COVID-19 has raised the possibility that the lethality related to infection with the SARS-CoV-2 is possibly related to an uncontrolled autoimmune response induced by the virus (*106*), and the presence of auto-antibodies against type I IFNs in patients with life-threatening COVID-19 has now been confirmed (*107*).

In conclusion, screening of our library of organic dyes and related novel drug-like compounds led to the identification of several small-molecule compounds showing promising broad-spectrum inhibition of the PPI between coronavirus spike proteins and their cognate ACE2 receptor. For several of them, including dyes, such as Congo red and direct violet 1, but especially novel non-dye compounds, such as DRI-C23041, we have confirmed that they are able to inhibit the entry of a SARS-CoV-2-S expressing pseudovirus into ACE2-expressing cells in a concentration-dependent manner. While activities might require further optimization, these results provide clear proof-of-principle evidence that this PPI, critical for CoV attachment and entry, is susceptible to small-molecule inhibition, making it feasible to pursue such alternative therapeutic options for the prevention and treatment of COVID-19 as oral or inhaled medications.

## MATERIALS AND METHODS

Commercial grade reagents and solvents were purchased from VWR (Radnor, PA, USA) and Sigma-Aldrich (St. Louis, MO, USA) and directly used without further purification. Chemicals, reagents, and the overwhelming majority of compounds used here were obtained from Sigma-Aldrich (St. Louis, MO, USA) and used as such; purity values are available on the manufacturer website. Some organic dyes (e.g., acid brown M, direct violet 1, and chlorazol black BH) were from TCI America (Portland, OR, USA); direct red 80 was from Santa Cruz Biotechnology (Dallas, TX, USA); gallein, NF023, and suramin were from Tocris Bioscience (Biotechne, Minneapolis, MN, USA).

### Chemistry

#### General methods

All reactions were carried out in oven- or flame-dried glassware under an atmosphere of dry argon (unless otherwise noted) and were magnetically stirred and monitored by analytical thin-layer chromatography (TLC) using Merck (Kenilworth, NJ, USA) pre-coated silica gel plates with F_254_ indicator (except if otherwise indicated). Visualization was accomplished by UV light (256 nm) with a combination of potassium permanganate and/or vanillin solution as an indicator. Flash column chromatography was performed according to the method of Still (*108*) using silica gel 60 (mesh 230-400; EMD Millipore, Billerica, MA, USA).

All newly synthesized compounds were characterized with ^1^H NMR, ^13^C NMR, high-resolution mass spectrometry (HRMS), and infrared (IR) spectroscopy – detailed data are provided in the Supplementary Material. Chemical shifts are reported in ppm relative to TMS. DMSO-*d*_6_ (2.50 ppm) was used as a solvent for ^1^H NMR and ^13^C NMR. ^1^H NMR and ^13^C NMR spectra were recorded on Bruker Avance 300 (300 MHz ^1^H), 400 (400 MHz ^1^H, 100 MHz ^13^C), and 500 (500 MHz ^1^H, 125 MHz ^13^C). Chemical shift values (*δ*) are reported in ppm relative to Me_4_Si (*δ* 0.0 ppm) unless otherwise noted. Proton spectra are reported as *δ* (multiplicity, coupling constant *J*, number of protons). Multiplicities are indicated by *s* (singlet), *d* (doublet), *t* (triplet), *q* (quartet), *p* (quintet), *h* (septet), *m* (multiplet), and *br* (broad). IR spectra were recorded with a FT-IR spectrophotometer Paragon 1000 (PerkinElmer). Mass spectra were obtained at the Mass Spectrometry Research and Education Center, Department of Chemistry, University of Florida (Gainesville, FL, USA). Low-resolution ES (electron spray) mass spectra were carried out with Finnigan LCQ DECA/Agilent 1100 LC/MS mass spectrometer (Thermo Fisher Scientific, Waltham, MA, USA). High-resolution mass spectra were recorded on an Agilent 6220 ESI TOF (Santa Clara, CA, USA) mass spectrometer. Analysis of sample purity was performed on an Agilent (Palo Alto, CA, USA) 1100 series HPLC system with a Thermoscientific Hypurity C8 (5 μm; 2.1 × 100 mm + guard column). HPLC conditions were as follows: solvent A = water with 2 mM ammonium acetate, solvent B = methanol with 2 mM ammonium acetate, and flow rate = 0.2 mL/min. Compounds were eluted with a gradient of A/B = 80:20 at 0 min to 0:100 at 50 min. Purity was determined via integration of UV spectra at 254 nm, and all tested compounds have a purity of ≥ 95%. All synthesized target compounds were tested as triethylamine salts unless otherwise stated. Details of the synthesis and structure conformation for all DRI-C compounds used here are summarized in the Supplementary Material (Supplementary Methods and Supplementary Schemes S1-S6).

### Binding Assays

SARS-CoV-2 S1 and RBD (cat. no. 40591-V08H and 40592-V08H), SARS-CoV S1+S2 (cat. no. 40634-V08B), HCoV-NL63 S1 (cat. no. 40600-V08H; all with His tag) and ACE2-Fc (cat. no. 10108-H05H) used in the binding assay were obtained from SinoBiological (Wayne, PA, USA). The TNF-R1:Fc receptor (cat. no. ALX-522-013-C050) and its FLAG-tagged TNF-α ligand (cat. no. ALX-522-008-C050) were obtained from Enzo Life Sciences (San Diego, CA, USA). Binding inhibition assays were performed in a 96-well cell-free format similar to the one described before (*53, 56–58*). Briefly, microtiter plates (Nunc F Maxisorp, 96-well; Thermo Fisher Scientific, Waltham, MA, USA) were coated overnight at 4 °C with 100 μL/well of Fc-conjugated ACE2 receptor diluted in PBS pH 7.2. This was followed by blocking with 200 μL/well of SuperBlock (PBS) (Thermo Fisher Scientific) for 1 h at RT (*60*). Then, plates were washed twice using washing solution (PBS pH 7.4, 0.05% Tween-20) and tapped dry before the addition of the tagged ligand (SARS-CoV-2 S1 or RBD) and test compounds diluted in binding buffer (100 mM HEPES, pH 7.2) to give a total volume of 100 μL/well. After 1 h incubation, three washes were conducted, and a further 1 h incubation with anti-His HRP conjugate (BioLegend; San Diego, CA, USA; cat. no. 652504) diluted (1:2500) in SuperBlock (PBS) was used to detect the bound His-tagged ligand. Plates were washed four times before the addition of 100 μL/well of HRP substrate TMB (3,3′,5,5′-tetramethylbenzidine) and kept in the dark for up to 15 min. The reaction was stopped using 20 μL of 1M H_2_SO_4_, and the absorbance value was read at 450 nm. The plated concentrations of ACE2 receptor and corresponding concentrations of the ligand used in the inhibitory assays were: 1.0 μg/mL ACE2 with 0.5 μg/mL SARS-CoV-2 RBD, 2.0 μg/mL ACE2 with 1.5 μg/mL SARS-CoV-2 S1, and 1.0 μg/mL ACE2 with 1 μg/mL SARS-CoV S1S2.. These values were selected following preliminary testing to optimize response (i.e., to produce a high-enough signal at conditions close to half-maximal response, EC_50_). Binding assessments for TNF-R1–TNF-α were performed as previously described using TNF-R1 at 0.3 μg/mL and TNF-α at 0.02 μg/mL (*59*), with the exception of using SuperBlock as the blocking buffer here. Stock solutions of compounds at 10 mM in DMSO were used.

### Protein Thermal Shift (Differential Scanning Fluorimetry)

This assay was used as described before (*59*) and following standard protocols from literature (*78, 79*) to establish which protein binds our compounds. SYPRO Orange (ThermoFisher; Waltham, MA, USA) was used as the fluorescence detection dye with an RT-PCR machine (StepOnePlus, Applied Biosystems, Foster City, CA, USA; detection on ROX channel, 575/602 nm) programmed to equilibrate samples at 25 °C for 90 s and then increase temperature to 99 °C by 0.4 °C every 24 s before taking a reading. Melting point of the protein is considered the lowest point of the first derivative plot, as calculated by the software included with the RT-PCR machine. Optimal concentrations were determined by performing a series of preliminary scans at various concentrations of protein, compound, and dye (SARS-CoV-2-RBD 0.05 mg/mL, hACE2-Fc 0.05 mg/mL, SYPRO Orange 4×, 100 mM HEPES buffer, 10 μM of test compound).

### SARS-CoV-2 Pseudovirus Assay

Fluorescent biosensors from Montana Molecular (Bozeman, MT, USA; cat. no. C1100R and C1100G) were used per the instructions of the manufacturer with minor modifications. Briefly, HEK293T cells (ATCC, Manassas, VA, USA; cat. no. CRL-1573) were seeded onto 96-well plates at a density of 50,000 cells per well in 100 μL complete medium (DMEM supplemented with 10% fetal bovine serum). A transduction mixture containing ACE2 BacMan Red-Reporter virus (1.8×10^8^ Vg/mL) and 2 mM sodium butyrate prepared in complete medium was added (50 μL per well) and incubated for 24 h at 37°C and 5% CO_2_. Medium was removed, washed once with PBS, and replaced with 100 μL fresh medium containing the compound under study, pre-incubating for 30 min at 37°C and 5% CO_2_. A transduction mixture containing Pseudo SARS-CoV-2 Green-Reporter pseudovirus (3.3×10^8^ Vg/mL) and 2 mM sodium butyrate prepared in complete medium was added (50 μL per well) and incubated for 48 h at 37°C and 5% CO_2_. Medium was removed, washed once with PBS, replaced with 150 μL fresh medium, and cells incubated for additional 48 h at 37°C and 5% CO_2_. Cell fluorescence was detected using an EVOS FL microscope (Life Technologies, Carlsbad, CA, USA) and quantified using the Analyze Particles tool after thresholding for the corresponding colors in ImageJ (US National Institutes of Health, Bethesda, MD, USA; (*109*)).

### Cytotoxicity Assay

For the MTS assay, HEK293T cells were cultured and prepared in the same manner as for the pseudovirus assay (up until the removal of test compounds there). Briefly, cells were added to a 96-well microtiter plate at a density of 50,000 cells/well in the absence or presence of various concentrations of compounds diluted in the same media. The plate was incubated at 37°C for 48 h. After washing three times with culture media, 20 μL per well of MTS, 3-(4,5-dimethylthiazol-2-yl)-5-(3-carboxymethoxyphenyl)-2-(4-sulfophenyl)-2H-tetrazolium, (Promega, Madison, WI, USA) was added to the plate at a final volume of 200 μL, and cells were incubated at 37°C for 2 h. Formazan levels were measured using a plate reader at 490 nm.

### Statistics and Data Fitting

All binding inhibition assays were performed as at least duplicates per plates, and all results shown are average of at least two independent experiments. As before (*56–58*), binding data were converted to percent inhibition and fitted with standard log inhibitor vs. normalized response models (*63*) using nonlinear regression in GraphPad Prism (GraphPad, La Jolla, CA, USA) to establish half-maximal effective or inhibitory concentrations (EC_50_, IC_50_).

## ABBREVIATIONS

ACE2: angiotensin converting enzyme 2
CoV: coronavirus
PPI: protein-protein interaction
SARS: severe acute respiratory syndrome
SMI: small-molecule inhibitor

## ACKNOWLEDGEMENTS

Financial support by the Diabetes Research Institute Foundation (www.diabetesresearch.org) is gratefully acknowledged. We thank the Mass Spectrometry Research and Education Center at the University of Florida (supported by funding from NIH S10 OD021758-01A1) for their prompt and professional service.

## AUTHOR CONTRIBUTIONS

DB performed most of the experiments, OA the pseudovirus assays, and JC the chemical synthesis. PB originated and designed the project, provided study guidance, analyzed the data, and wrote the draft manuscript. All authors contributed to writing and read the final manuscript.

## SUPPLEMENTARY MATERIALS

Supplementary material for this article is available at the end of this file and includes Supplementary Methods, Schemes S1 to S6, and Figures S1 to S3.

## SUPPLEMENTARY METHODS

### 8-(4′-Nitrobiphenyl-4-ylcarboxamido)naphthalene-1-sulfonic acid (16) and the general procedure for coupling

4′-Nitro[1,1′-biphenyl]-4-carboxylic acid (**14**) was synthesized by two steps as described in the literature (*110*). As described before (*58*), for the synthesis of **16** and as a general procedure of coupling a modified version of the coupling reaction from reference (*61*) was used (Scheme S1). Under an argon atmosphere, trimethylamine (1.74 mL, 12.5 mmol) was added dropwise to a mixture of 4′-nitro[1,1′-biphenyl]-4-carboxylic acid (**14**) (1.63 g, 6.7 mmol), *O*-(6-chlorobenzotriazol-1-yl)-*N*,*N*,*N*′,*N*′-tetramethyluronium hexafluorophosphate HCTU (2.7 g, 6.5 mmol) and DMF (10 mL) at 0 °C and the resulting reaction mixture was stirred for 1 h at the same temperature. Subsequently, 8-amino-1-naphthalenesulfonic acid **15** (1.50 g, 6.7 mmol) was added at the same temperature. The resulting reaction mixture was allowed to stir overnight at room temperature (RT). Diethyl ether (50 mL) was added to the reaction mixture, and a yellow precipitate formed. This precipitate was collected by filtration and washed with diethyl ether (3 × 10 mL) to afford the triethylamine salt of **16** as a yellow solid (2.4 g, 66%). ^1^H-NMR (500 MHz, DMSO-*d*_6_): *δ* 12.70 (s, 1H), 8.79 (br, 1H), 8.38 - 8.29 (m, 5H), 8.20 (d, *J* = 7.6 Hz, 1H), 8.09 (d, *J* = 8.8 Hz, 2H), 8.02 (d, *J* = 8.1 Hz, 1H), 7.95 (d, *J* = 8.4 Hz, 2H), 7.83 (d, *J* = 7.8 Hz, 1H), 7.59 (t, *J* = 7.9 Hz, 1H), 7.48 (t, *J* = 7.6 Hz, 1H), 3.08 (q, *J* = 7.2 Hz, 6H), 1.15 (t, *J* = 7.3 Hz, 9H); ^13^C-NMR (125 MHz, DMSO-*d*_6_): δ 164.8, 147.0, 145.8, 141.8, 140.1, 135.9, 135.8, 133.2, 131.9, 128.9, 128.1, 127.4, 127.0, 126.1, 125.3, 124.2, 124.1, 124.0, 123.0, 45.8, 8.6; FTIR (neat) ν_max_ 3028, 2735, 1668, 1596, 1515, 1494, 1480, 1429, 1393, 1339, 1328, 1280, 1231, 1186, 1162, 1126, 1110, 1039, 1010, 924, 894, 869, 854, 843, 826, 788, 763, 740, 692 cm^−1^; HRMS (ESI) [M + H]^+^ calcd. for C_23_H_17_N_2_O_6_S^+^, 449.0802; found, 449.0781.

### 8-(4′-Aminobiphenyl-4-ylcarboxamido)naphthalene-1-sulfonic acid (17) and the general procedure for hydrogenation

For the synthesis of **17** and as a general procedure of hydrogenation a modified version of the corresponding reaction from reference (*62*) was used as described before (*58*) (Scheme S1). A mixture of 8-(4′-nitrobiphenyl-4-ylcarboxamido)naphthalene-1-sulfonic acid (**16**) (2.8 g, 5.1 mmol) and 10% Pd on carbon (81 mg) in a solvent mixture of EtOH (6 mL) and DMF (3 mL) was hydrogenated (H_2_ balloon) at 80 °C for 3.5 h. The reaction mixture was filtered via a short pad of Celite®, concentrated *in vacuo*, and recrystallized from MeOH to afford the triethylamine salt of **17** as a white solid (2.3 g, 86%). ^1^H-NMR (500 MHz, DMSO-*d*_6_): *δ* 12.56 (s, 1H), 8.80 (br, 1H), 8.31 (dd, *J* = 1.1, 6.1 Hz, 1H), 8.25–8.12 (m, 3H), 8.00 (d, *J* = 7.2 Hz, 1H), 7.80 (d, *J* = 7.3 Hz, 1H), 7.67 (d, *J* = 8.3 Hz, 2H), 7.57 (t, *J* = 7.8 Hz, 1H), 7.49 (d, *J* = 7.0 Hz, 2H), 7.47 (t, *J* = 7.7 Hz, 1H), 6.68 (d, *J* = 8.4 Hz, 2H), 5.33 (s, 2H), 3.05 (q, *J* = 7.2 Hz, 6H), 1.14 (t, *J* = 7.3 Hz, 9H); ^13^C-NMR (125 MHz, DMSO-*d*_6_): δ 165.2, 148.9, 143.1, 141.8, 135.7, 133.4, 132.5, 131.8, 128.6, 127.4, 127.3, 126.3, 125.7, 125.3, 124.6, 124.1, 123.9, 123.0, 114.2, 45.7, 8.6; FTIR (neat) ν_max_ 3431, 3338, 3227, 3006, 2712, 1648, 1602, 1531, 1492, 1474, 1429, 1397, 1331, 1284, 1227, 1196, 1184, 1161, 1061, 1036, 1008, 921, 891, 823, 788, 761, 725, 704, 663 cm^−1^; HRMS (ESI) [M + H]^+^ calcd. for C_23_H_19_N_2_O_4_S^+^, 419.1060; found, 419.1058.

### 8-(4′-(4-Nitrobenzamido)biphenyl-4-ylcarboxamido)naphthalene-1-sulfonic acid (7, DRI-C21041)

Preparation of compound **7** followed the synthetic scheme of Scheme S1 via intermediaries **16** and **17**. The general procedure for coupling as described for **16** was followed with 4-nitrobenzoic acid (**18**) (0.50 g, 3.0 mmol) and 8-(4′-aminobiphenyl-4-ylcarboxamido)naphthalene-1-sulfonic acid (**19**) (1.20 g, 2.3 mmol) to give the triethylamine salt of the title compound **7** as a yellow solid (1.35 g, 87%) (99% pure by HPLC analysis (UV spectra at 254 nm)). ^1^H-NMR (500 MHz, DMSO-*d*_6_): *δ* 12.64 (s, 1H), 10.75 (s, 1H), 8.85 (br, 1H), 8.39 (d, *J* = 8.1 Hz, 2H), 8.36–8.16 (m, 6H), 8.02 (d, *J* = 8.2 Hz, 1H), 7.95 (d, *J* = 8.2 Hz, 2H), 7.91–7.81 (m, 5H), 7.59 (t, *J* = 7.7 Hz, 1H), 7.49 (t, *J* = 7.5 Hz, 1H), 3.07 (q, *J* = 7.2 Hz, 6H), 1.15 (t, *J* = 7.1 Hz, 9H); ^13^C-NMR (125 MHz, DMSO-*d*_6_): δ 165.1, 164.1, 149.2, 142.0, 141.8, 140.6, 138.8, 135.8, 134.9, 134.2, 133.3, 132.0, 129.4, 128.8, 127.5, 127.3, 126.0, 125.9, 125.4, 124.4, 124.1, 123.7, 123.0, 120.8, 45.7, 8.7; FTIR (neat) ν_max_ 3360, 3017, 2714, 1679, 1666, 1592, 1521, 1489, 1432, 1416, 1398, 1340, 1321, 1279, 1235, 1194, 1152, 1131, 1102, 1038, 1009, 929, 895, 864, 852, 824, 761, 708, 675, 661 cm^−1^; HRMS (ESI) [M − H]^−^ calcd. for C_30_H_20_N_3_O_7_S^−^, 566.1027; found, 566.1054.

### 5-(4’-(4-Nitrobenzamido)biphenyl-4-ylcarboxamido)naphthalene-2-sulfonic acid (8, DRI-C23041)

Preparation of compound **8** followed the synthetic scheme of Scheme S2. The general procedure for the coupling reaction as described earlier was followed with 4’-(4-nitrobenzamido)biphenyl-4-carboxylic acid **20** (181 mg, 0.5 mmol) and 5-aminonaphthalene-2-sulfonic acid **21** (112 mg, 0.5 mmol) to give the triethylamine salt of the title compound as a yellowish solid (85 mg, 30%) (>99% pure by HPLC analysis (UV spectra at 254 nm)). ^1^H NMR (500 MHz, DMSO-*d*_6_): *δ* 10.73 (s, 1H), 10.49 (s, 1H), 8.86 (br, 1H), 8.39 (d, *J* = 8.4 Hz, 2H), 8.30 - 8.15 (m, 5H), 8.05 - 7.88 (m, 6H), 7.84 (d, *J* = 8.4 Hz, 2H), 7.77 (d, *J* = 8.8 Hz, 1H), 7.65 (d, *J* = 7.1 Hz, 1H), 7.58 (t, *J* = 7.9 Hz, 1H), 3.08 (q, *J* = 6.8 Hz, 6H), 1.16 (t, *J* = 7.2 Hz, 9H); ^13^C NMR (125 MHz, DMSO-*d*_6_): δ 165.8, 164.0, 149.2, 145.6, 142.6, 140.5, 138.9, 134.7, 133.8, 133.0, 132.9, 129.3, 128.9, 128.6, 127.2, 126.9, 126.2, 126.0, 124.4, 123.9, 123.6, 123.1, 120.9, 45.8, 8.6; HRMS (ESI) [M-H]^−^ calcd. for C_30_H_20_N_3_O_7_S^−^, 566.1027; found, 566.1052.

### 5-(4’-(4-Nitrobenzamido)biphenyl-4-ylcarboxamido)naphthalene-1-sulfonic acid (9, DRI-C24041)

Preparation of compound **9** followed the synthetic scheme of Scheme S2. The general procedure for the coupling reaction as described earlier was followed with 4’-(4-nitrobenzamido)biphenyl-4-carboxylic acid **20** (181 mg, 0.5 mmol) and 5-aminonaphthalene-1-sulfonic acid **22** (112 mg, 0.5 mmol) to give the triethylamine salt of the title compound as a yellowish solid (210 mg, 63%) (>99% pure by HPLC analysis (UV spectra at 254 nm)). ^1^H NMR (500 MHz, DMSO-*d*_6_): *δ* 10.73 (s, 1H), 10.49 (s, 1H), 8.86 (d, *J* = 7.9 Hz, 2H), 8.39 (d, *J* = 8.2 Hz, 2H), 8.24 (d, *J* = 8.4 Hz, 2H), 8.21 (d, *J* = 8.0 Hz, 2H), 8.02 (d, *J* = 7.6 Hz, 2H), 7.97 (d, *J* = 8.3 Hz, 2H), 7.90 (d, *J* = 8.0 Hz, 2H), 7.85 (d, *J* = 8.3 Hz, 2H), 7.59 (d, *J* = 7.1 Hz, 1H), 7.56 (d, *J* = 7.2 Hz, 1H), 7.49 (t, *J* = 7.9 Hz, 1H), 3.07 (q, *J* = 7.3 Hz, 6H), 1.16 (t, *J* = 7.4 Hz, 9H); ^13^C NMR (125 MHz, DMSO-*d*_6_): δ 165.8, 164.0, 149.2, 144.3, 142.5, 140.5, 138.8, 134.6, 133.7, 132.9, 130.0, 129.8, 129.3, 128.5, 127.2, 126.3, 126.2, 125.0, 124.8, 124.53, 124.46, 123.9, 123.6, 120.8, 45.8, 8.6; HRMS (ESI) [M-H]^−^ calcd. for C_30_H_20_N_3_O_7_S^−^, 566.1027; found, 566.1025.

### 8-(4-(5-(4-Nitrobenzamido)pyridin-2-yl)benzamido)naphthalene-1-sulfonic acid (11, DRI-C61041)

Preparation of compound **11** followed the synthetic scheme of Scheme S3. The general procedure for the coupling reaction as described earlier was followed with 4-nitrobenzoic acid **18** (109 mg, 0.65 mmol) and the triethyl amine salt of 8-(4-(5-aminopyridin-2-yl)benzamido)naphthalene-1-sulfonic acid **28** (260 mg, 0.50 mmol) to give the triethylamine salt of the title compound as a yellowish solid (260 mg, 78%) (>99% pure by HPLC analysis (UV spectra at 254 nm)). ^1^H NMR (500 MHz, DMSO-*d*_6_): *δ* 12.67 (s, 1H), 10.89 (s, 1H), 9.09 (s, 1H), 8.80 (br, 1H), 8.38 (d, *J* = 8.3 Hz, 2H), 8.36 (d, *J* = 8.1 Hz, 1H), 8.32 (d, *J* = 7.1 Hz, 1H), 8.28 (d, *J* = 8.0 Hz, 2H), 8.24 (d, *J* = 8.4 Hz, 2H), 8.21 (d, *J* = 8.1 Hz, 3H), 8.14 (d, *J* = 8.5 Hz, 1H), 8.01 (d, *J* = 8.0 Hz, 1H), 7.82 (d, *J* = 8.0 Hz, 1H), 7.58 (t, *J* = 7.8 Hz, 1H), 7.48 (t, *J* = 7.5 Hz, 1H), 3.07 (q, *J* = 6.1 Hz, 6H), 1.15 (t, *J* = 7.2 Hz 9H); ^13^C NMR (125 MHz, DMSO-*d*_6_): δ 165.1, 164.3, 150.6, 149.3, 141.8, 141.7, 140.5, 139.9, 135.8, 135.5, 134.9, 133.2, 131.9, 129.3, 128.5, 128.2, 127.5, 126.0, 125.7, 125.4, 124.3, 124.0, 123.6, 123.0, 120.5, 45.8, 8.6; HRMS (ESI) [M-H]^−^ calcd. for C_29_H_20_N_4_O_7_S^−^, 567.0980; found, 567.0998.

### 8-(4-(5-(4-Nitrobenzamido)thiophen-2-yl)benzamido)naphthalene-1-sulfonic acid (12, DRI-C71041)

Preparation of compound **12** followed the synthetic scheme of Scheme S4. The general procedure for the coupling reaction as described earlier was followed with 4-nitrobenzoyl chloride **34** (102 mg, 0.6 mmol) and the triethyl amine salt of 8-(4-(5-aminothiophen-2-yl)benzamido)naphthalene-1-sulfonic acid **33** (212g, 0.5 mmol) to give the triethylamine salt of the title compound as a red solid (140 mg, 42%) (>99% pure by HPLC analysis (UV spectra at 254 nm)). ^1^H NMR (500 MHz, DMSO-*d*_6_): *δ* 12.59 (s, 1H), 12.05 (s, 1H), 8.40 (d, *J* = 8.4 Hz, 2H), 8.32 (d, *J* = 7.0 Hz, 1H), 8.27 (d, *J* = 8.5 Hz, 2H), 8.20 (d, *J* = 7.3 Hz, 3H), 8.02 (d, *J* = 8.0 Hz, 1H), 7.82 (d, *J* = 7.8 Hz, 1H), 7.76 (d, *J* = 8.0 Hz, 2H), 7.58 (t, *J* = 7.8 Hz, 1H), 7.52 (d, *J* = 3.6 Hz, 1H), 7.48 (t, *J* = 7.7 Hz, 1H), 7.04 (d, *J* = 3.5 Hz, 1H); ^13^C NMR (125 MHz, DMSO-*d*_6_): δ 164.9, 161.6, 149.4, 141.8, 140.1, 138.5, 136.7, 135.8, 133.59, 133.57, 133.3, 131.9, 129.3, 129.0, 127.4, 125.9, 125.3, 124.2, 124.1, 124.0, 123.7, 123.0, 122.3, 114.4; HRMS (ESI) [M-H]^−^ calcd. for C_28_H_18_N_3_O_7_S_2_^−^, 572.0592; found, 572.0611.

### 4-Hydroxy-8-(4’-(4-methoxycarbonyl)biphenyl-4-ylcarboxamido)naphthalene-2-sulfonic acid (10, DRI-C2204745)

Preparation of compound **10** followed the synthetic scheme of Scheme S5. The general procedure for the coupling reaction as described earlier was followed with methyl 4-(chlorocarbonyl)benzoate **39** (119 mg, 0.6 mmol) and **38** (240 mg, 0.5 mmol) to give the title compound as a white solid (160 mg, 54%) (>99% pure by HPLC analysis (UV spectra at 254 nm)). 1H NMR (500 MHz, DMSO-*d*_6_): *δ* 10.61 (s, 1H), 10.48 (s, 1H), 10.28 (s, 1H), 8.19 (d, *J* = 8.0 Hz, 2H), 8.14 (d, *J* = 8.6 Hz, 2H), 8.12 (d, *J* = 8.5 Hz, 2H), 8.08 (d, *J* = 8.4 Hz, 1H), 7.97 (d, *J* = 8.4 Hz, 2H), 7.90 (d, *J* = 8.0 Hz, 2H), 7.89 – 7.77 (m, 3H), 7.56 (d, *J* = 7.1 Hz, 1H), 7.50 (t, *J* = 7.9 Hz, 1H), 7.23 (s, 1H), 3.91 (s, 3H); ^13^C NMR (125 MHz, DMSO-*d*_6_): δ 165.75, 165.73, 164.8, 152.9, 146.0, 142.6, 139.1, 139.0, 134.5, 134.4, 132.9, 132.1, 130.2, 129.2, 128.5, 128.2, 127.2, 126.2, 125.4, 125.1, 124.5, 120.8, 120.4, 110.6, 106.3, 52.5; HRMS (ESI) [M-H]^−^ calcd. for C_32_H_23_N_2_O_8_S^−^, 595.1181; found, 595.1199.

### N^4^,N^4^’-bis(2-sulfonic acid-4-hydroxynaphthalen-6-yl)biphenyl-4,4’-dicarboxamide (13, DRI-C91005)

Preparation of compound **13** followed the synthetic scheme of Scheme S6. 4,4’-Biphenyldicarbonyl chloride **40** (76.7 mg, 0.275 mmol) was added to the solution of 6-amino-4-hydroxynaphthalene-2-sulfonic acid **41** (120 mg, 0.5 mmol) in dioxane (2 mL) and water (2 mL) by portion at room temperature. During adding **40**, the pH value was kept within 4.0 to 5.0 by adding 1 N sodium carbonate dropwise. After reaction, the pH value was adjusted to 2. Dioxane and water were removed by high vacuum pump. The residue was transferred to a test tube and taken up with methanol at 80 °C. 2.0 mL of water was added. The reaction mixture was cooled to room temperature and filtered to give the title compound as a red solid (234 mg, 100%) (>99% pure by HPLC analysis (UV spectra at 254 nm)). ^1^H NMR (500 MHz, DMSO-*d*_6_): *δ* 10.49 (s, 2H), 10.11 (s, 2H), 8.63 (d, *J* = 2.0 Hz, 2H), 8.17 (d, *J* = 8.3 Hz, 4H), 7.98 (d, *J* = 8.3 Hz, 4H), 7.91 (dd, *J* = 8.9, 2.1 Hz, 2H), 7.85 (d, *J* = 8.9 Hz, 2H), 7.56 (s, 2H), 7.15 (d, *J* = 1.5 Hz, 2H); ^13^C NMR (100 MHz, DMSO-*d*_6_): δ 164.8, 163.0, 152.1, 150.5, 150.3, 145.0, 141.8, 136.0, 134.1, 130.7, 130.6, 128.3, 126.7, 124.5, 121.0; HRMS (ESI) [M-2H]^2–^ calcd. for C_34_H_22_N_2_O_10_S_2_^2−^,341.0352; found, 341.0361.

## SUPPLEMENTARY SCHEMES AND FIGURES

**Scheme S1.**
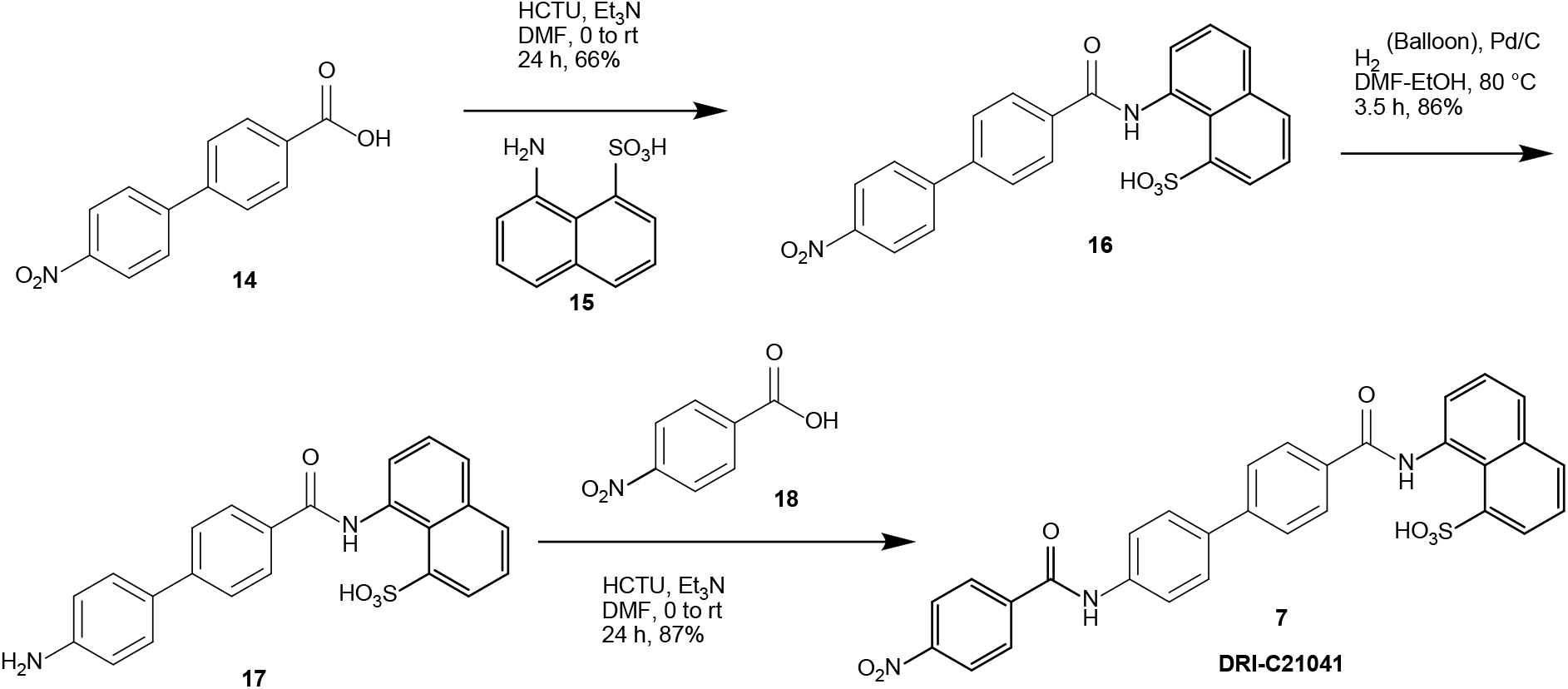
Synthesis of DRI-C21041 (**7**).

**Scheme S2.**
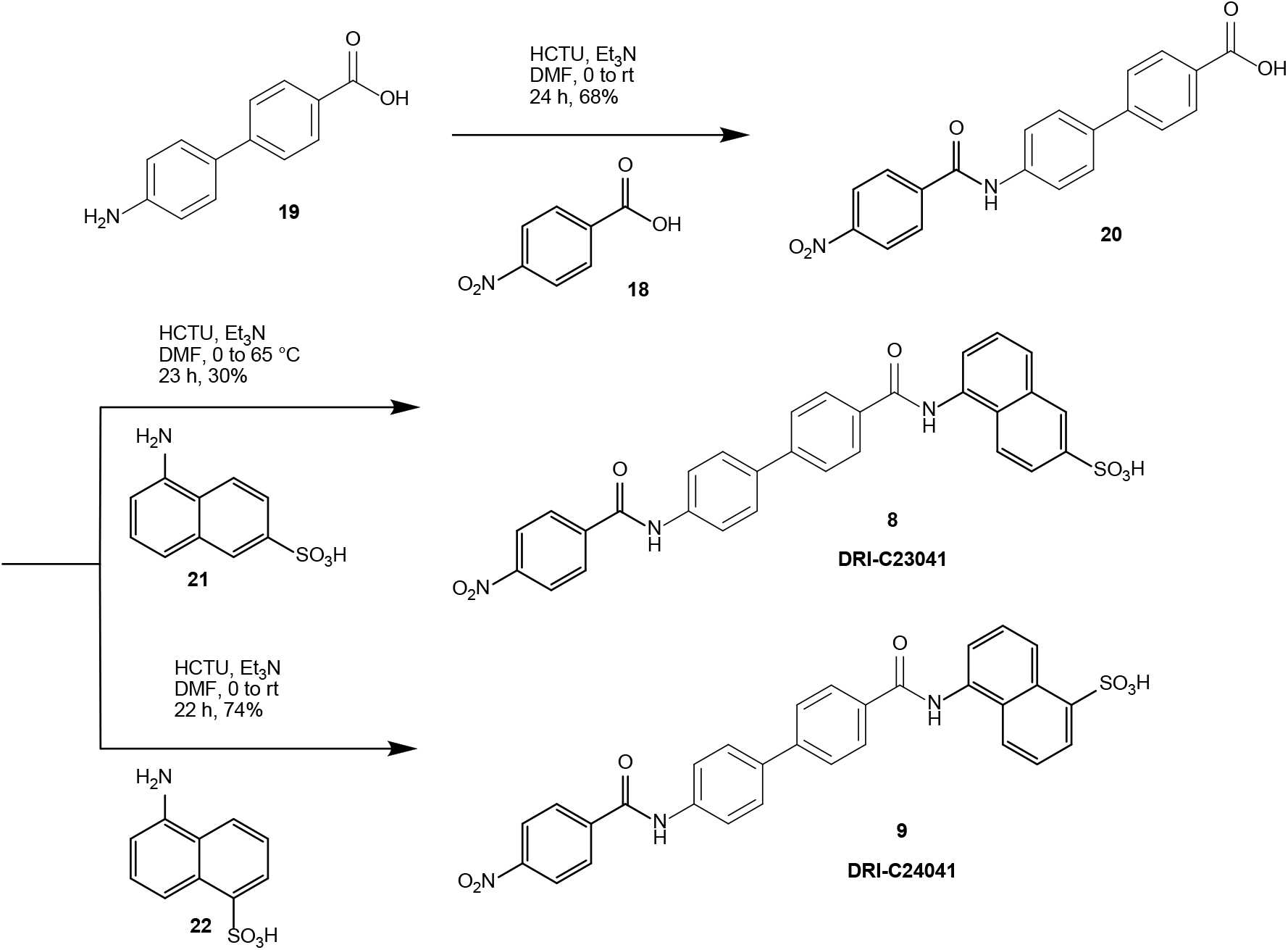
Synthesis of DRI-C23041 (**8**) and DRI-C24041 (**9**).

**Scheme S3.**
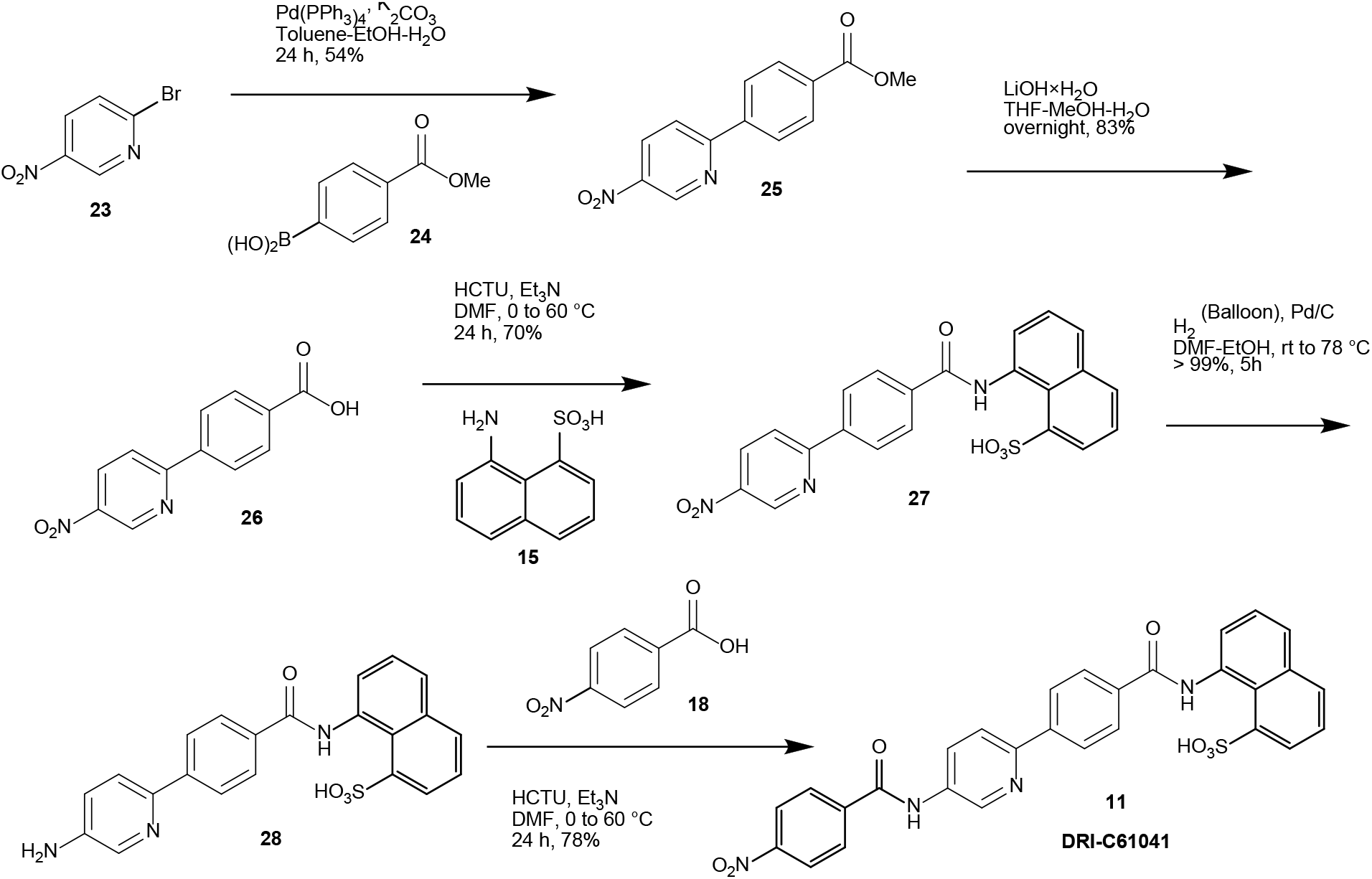
Synthesis of DRI-C61041 (**11**).

**Scheme S4.**
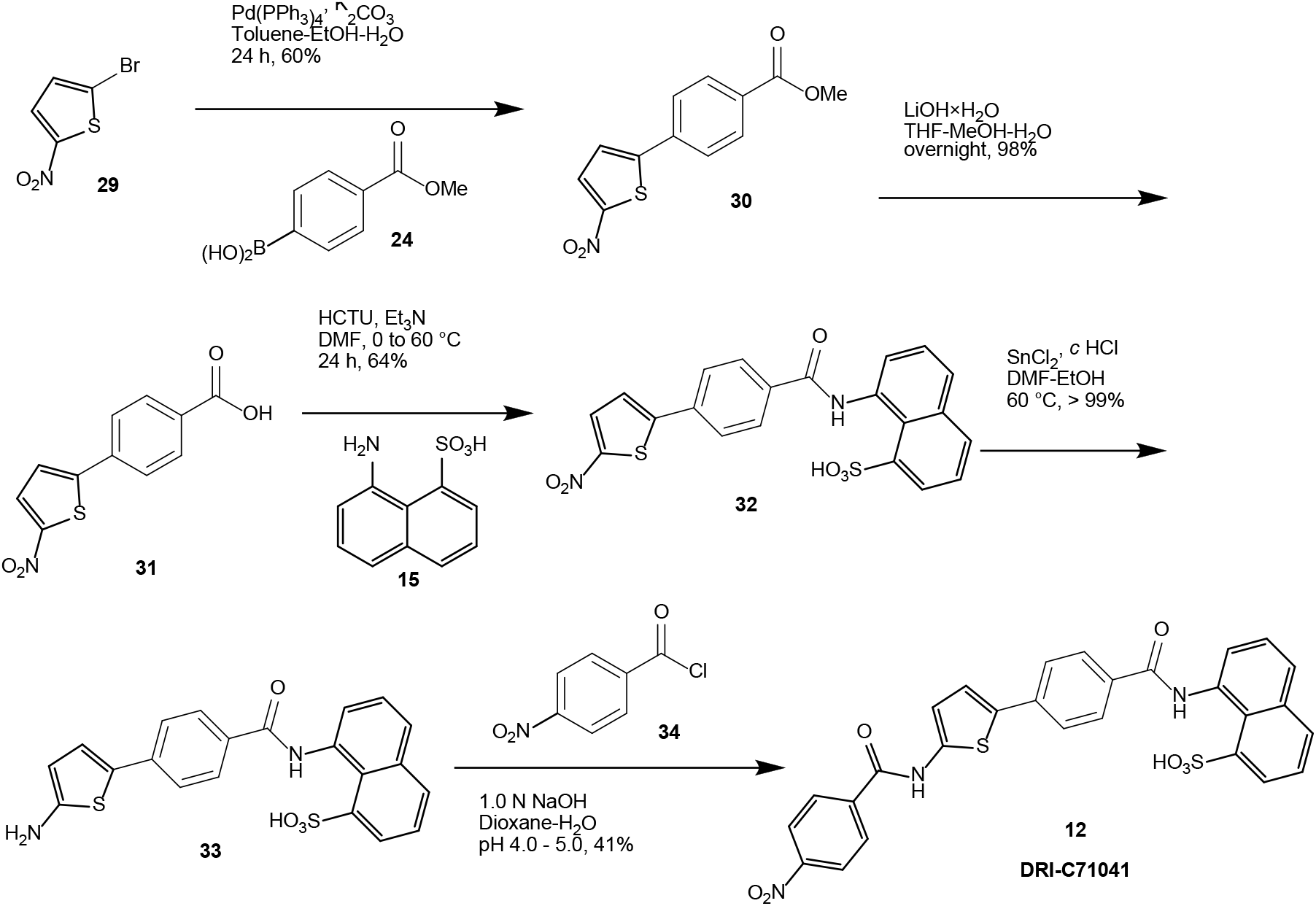
Synthesis of DRI-C71041 (**12**).

**Scheme S5.**
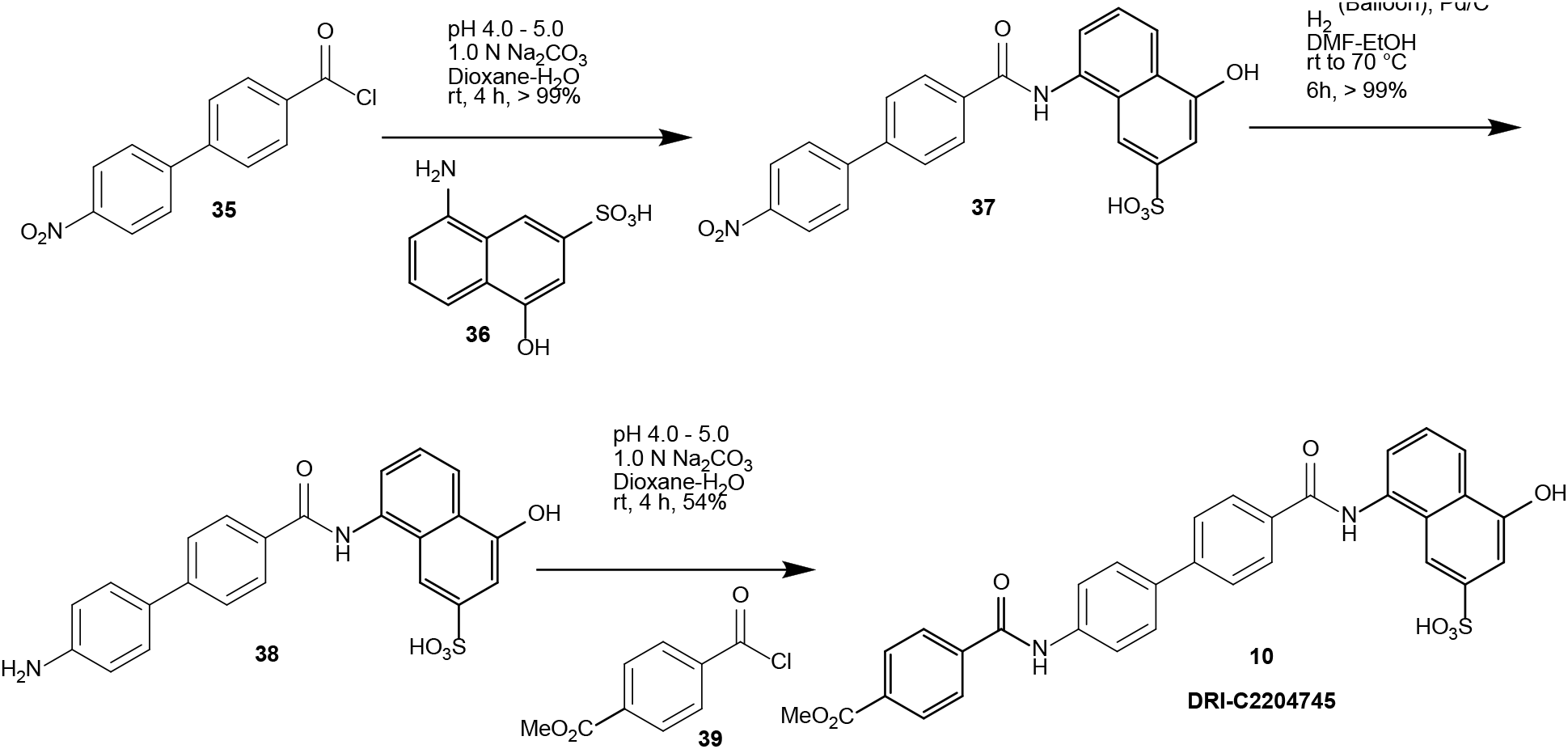
Synthesis of DRI-C2204745 (**10**).

**Scheme S6.**
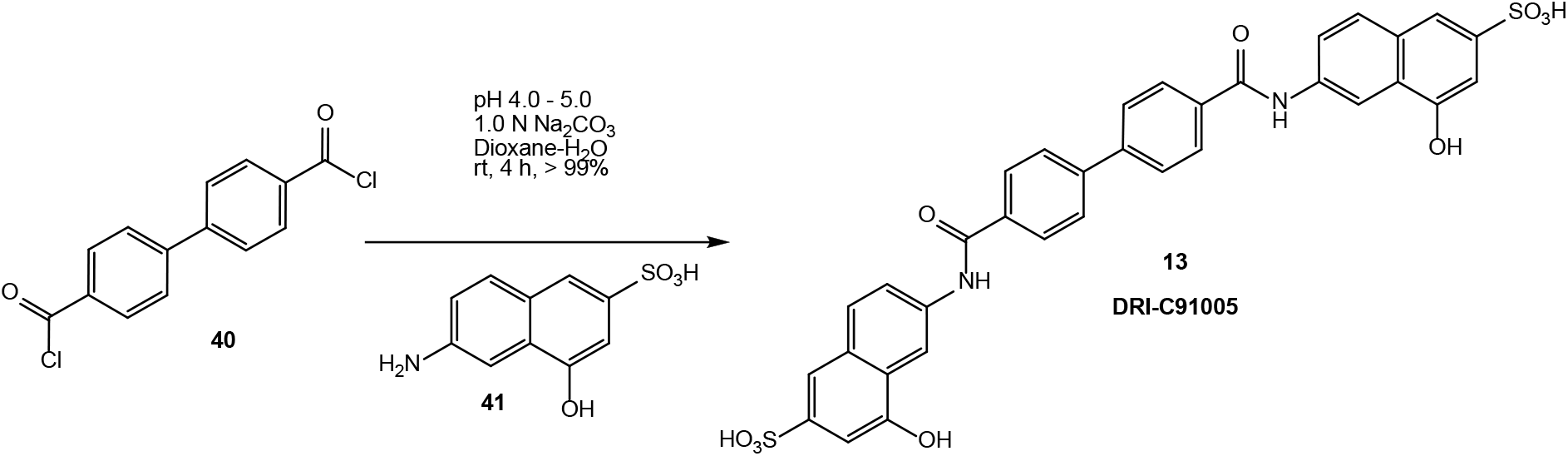
Synthesis of DRI-C91005 (**13**).

**Figure S1.**
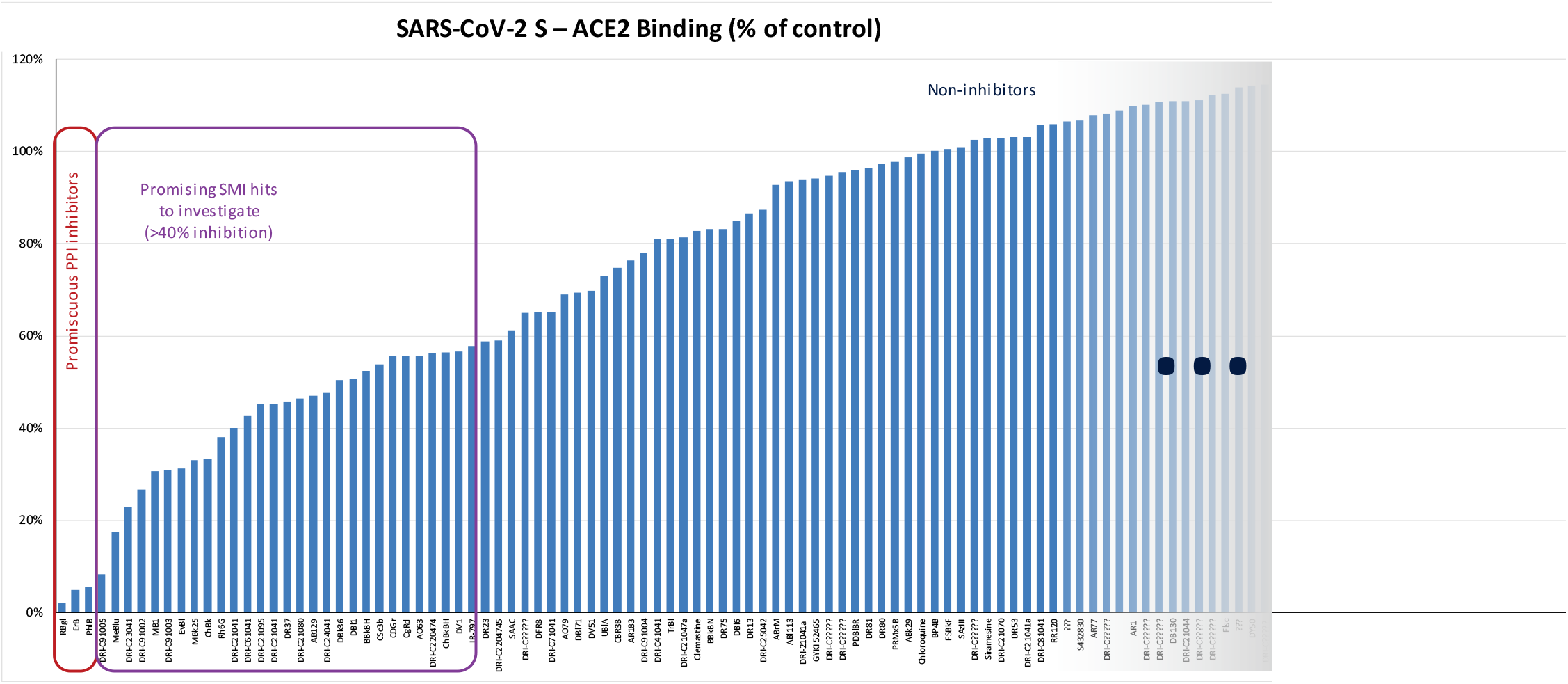
Inhibitory effect of selected compounds on SARS-CoV-2 RBD binding to hACE2 in our screening assay. Percent inhibition values obtained at 5 μM concentration shown normalized to control (100%). Rose Bengal, erythrosine B, and phloxine B that showed the highest activity are known promiscuous SMI of PPIs (*56*) and were included as positive controls. Compounds showing promising (>40%) inhibition were evaluated in detailed concentration-response assays to establish IC_50_ values.

**Figure S2.**
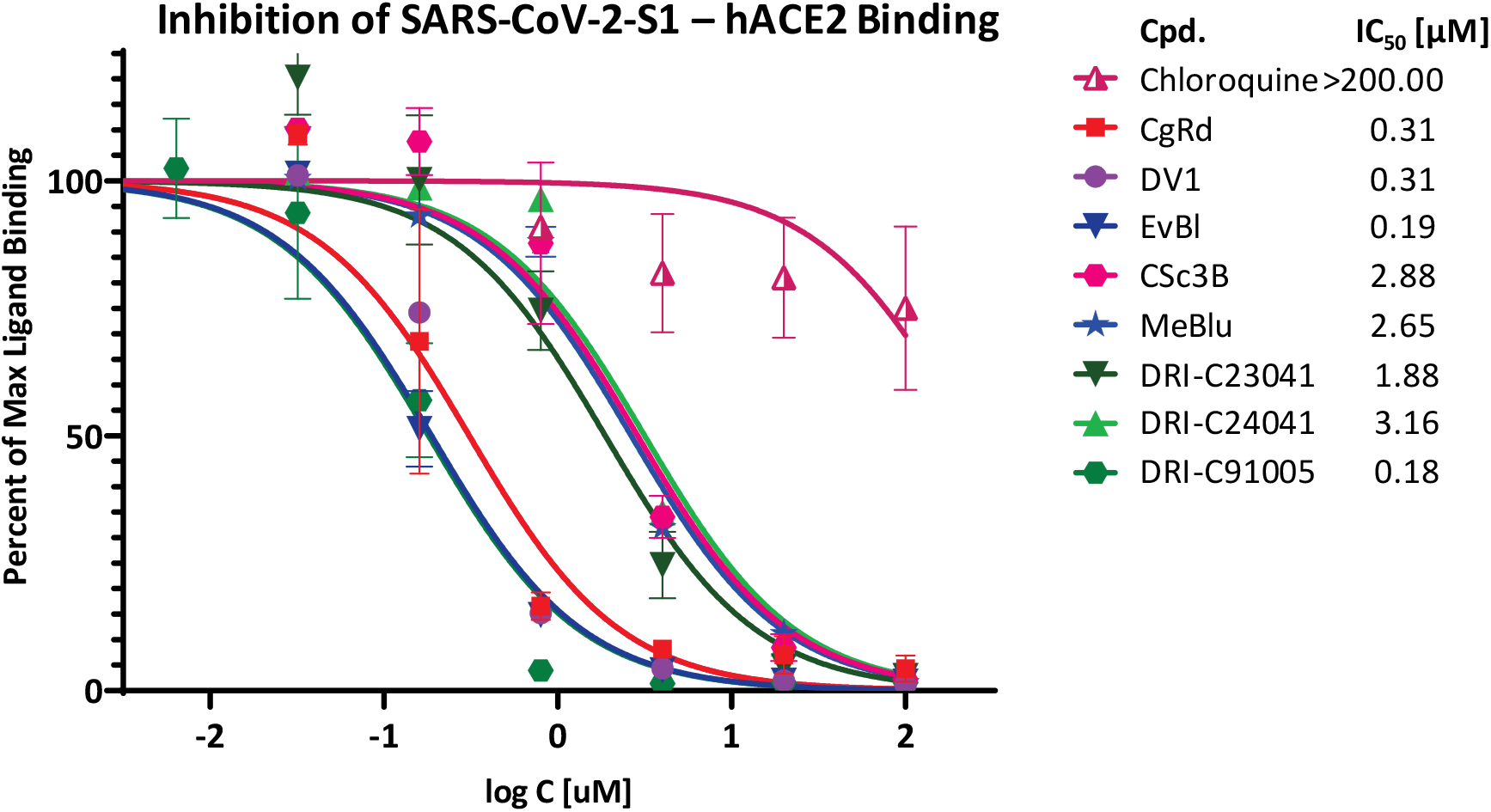
Concentration-dependent inhibition of SARS-CoV-2-S1 binding to ACE2 by compounds of the present study. Concentration-response curves obtained for the inhibition of the PPI between SARS-CoV-2-S1 (His-tagged, 1.5 μg/mL) and hACE2 (Fc-conjugated, 2 μg/mL) in cell-free ELISA-type assay with compounds tested. Data are mean ± SD from two experiments in duplicates and were fitted with standard sigmoid curves for IC_50_ determination as in Figure 3.

**Figure S3.**
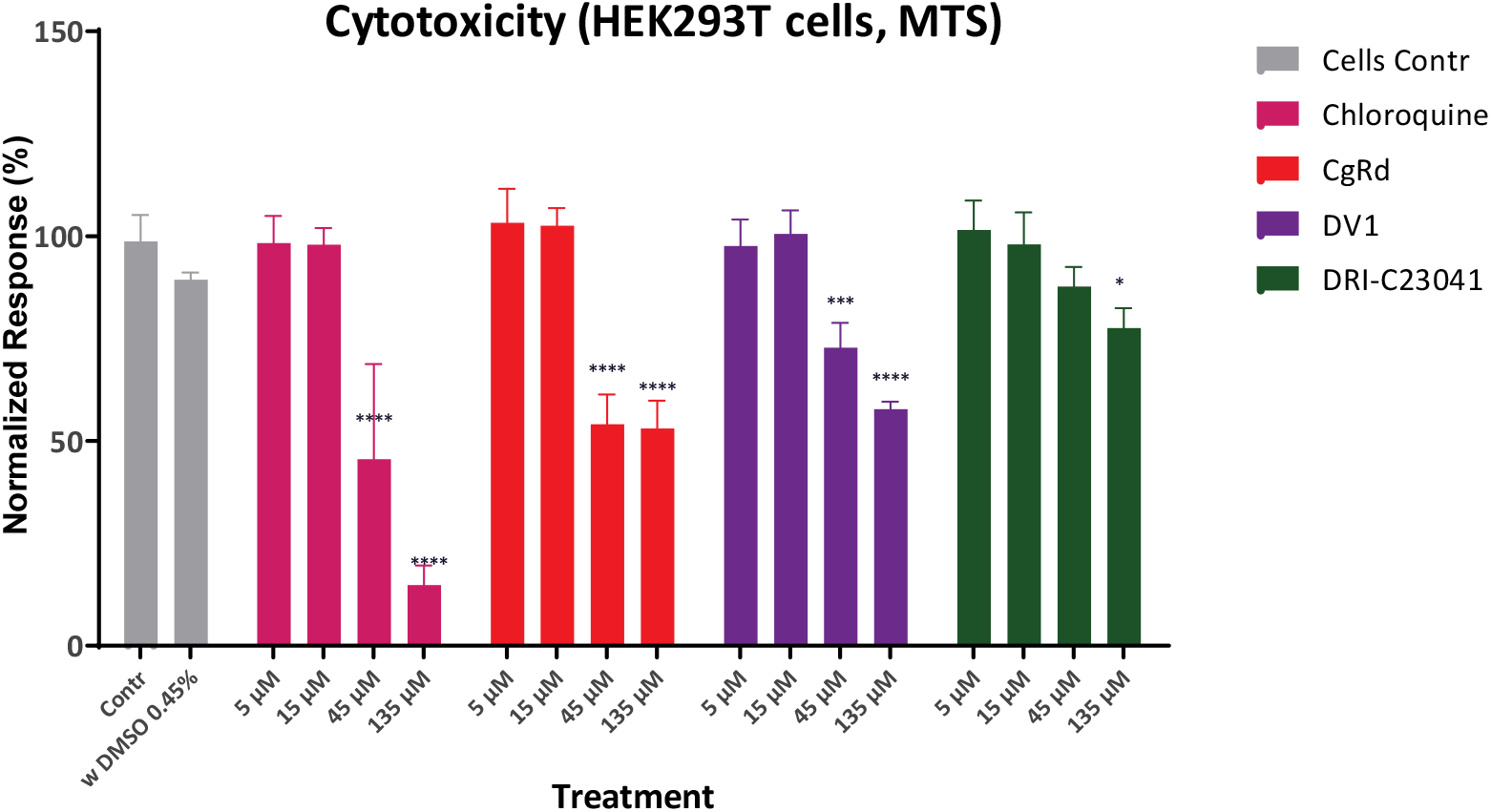
Toxicity assessment of selected compounds. using a standard MTS cytotoxicity assay in HEK293T cells. Data are average ± SD of two experiments in duplicate and were analyzed by one-way ANOVA followed by Dunnett’s multiple comparison test, ^****^*p* < 0.0001 vs. cells only, ^*^*p* < 0.05 vs. cells only.

